# Consciously detecting and recognizing a past visual word after its sensory trace is gone

**DOI:** 10.1101/2021.02.02.429359

**Authors:** Daphné Rimsky Robert, Matteo Lisi, Kevin Nguy, Roxane Jannin, Thomas Hardy, Nathan Beraud, Claire Sergent

## Abstract

What is the role of sensory processing in conscious perception? Current theories of consciousness are divided on this question. Some propose that conscious perception arises during the buildup of sensory representations. Other argue for a secondary process that broadcasts these representations to higher-level areas. This second view makes a counter-intuitive prediction: one could consciously perceive abstract representations untied to any low-level sensory feature. We tested this prediction by combining visual masking with retrospective cueing. We found that when visually masked words were followed by a semantically related auditory word, participants were better at detecting this past word and reporting its identity, but were strikingly unable to report its visual features (letter casing or position on screen). This suggests that retro-cueing can help a semantic representation reach awareness even after the associated sensory information has been masked. The mechanisms of conscious access might thus be largely independent from early sensory build-up.

## Introduction

When does the awareness of an external stimulus emerge during the course of its processing by the brain? This question stands as a significant juncture in ongoing debates about the neural mechanisms of conscious perception^1,2^. Three main processing stages can be distinguished, based on their neurophysiological and computational characteristics^3–6^. First, the onset of a new sensory input, visual, auditory or other, triggers a “feed-forward” sweep of activity, which, via ascending connections, hierarchically mobilizes the different areas in the sensory system, reaching the top levels within approximately 100ms^4,5^. In vision, it goes from the retina to the lateral geniculate nucleus, the primary visual cortex (V1), higher-level visual areas (V2, V3, V4…), and then parietal, temporal areas. At this stage, the encoding of the stimulus is typically quite faithful to the sensory input. But whenever activity reaches a specific level, this also triggers recurrent connections that modulate lower-levels in return (e.g. from V2 to V1). This can be identified as a “second stage” which allows the stabilization of sensory representations in a more interpreted form^3,7^. Finally, in a general “third stage”, functional connectivity extends beyond the sensory system, and typically involves frontal, parietal and cingulate areas, associated with attentional selection, opening a wide repertoire of cognitive functions^5,8,9^.

There is a large consensus in assuming that the first feed-forward sweep is an unconscious stage. Beyond that, the different theories diverge. Theories such as the Local Recurrent Processing^10^ or the Integrated Information Theory^11^ consider that conscious perception arises with the establishment of local recurrent loops within the sensory system^12^, notably because, at this stage, stimulus encoding departs from the strict rendering of the input to match the way the stimulus appears to the individual^13–15^. In this view, additional functional connectivity with supra-modal areas (third stage) only has a modulatory effect^16^.

In contrast, other views, such as the Global Neuronal Workspace^9,17^, or Higher-Order theories^18^, consider that this second stage is not sufficient to become conscious of the stimulus, and propose that conscious perception arises if and only if integrated sensory representations are further broadcast to a wider network of supra-modal areas (third stage), allowing a global and flexible functional impact on the cognition of the individual.

So far, experimental work contrasting brain activity for conscious versus non conscious perception has not been very conclusive: all processing stages have been found to correlate with conscious perception in one study or another^19–22^, including pre-stimulus activity^23,24^. These seemingly contradictory results reflect one limitation of this contrastive method, which reveals *everything* that correlates with the final perception of a stimulus, including events that might favor the future conscious perception of the stimulus, while not being directly part of the mechanisms of conscious perception.

To move forward, we must identify scenarios where various theories yield contrasted predictions, and subject them to testing. Here we highlight a strong and rather counter-intuitive prediction of high-level theories such as the Global Neuronal Workspace: if conscious access is largely independent from sensory build-up, then it should be possible to desynchronize conscious access from the timing of the external stimulus^25^. We have started to explore this possibility in several experiments testing whether conscious access to an initially missed stimulus could still be triggered by retrospective attention. We have shown that cueing spatial attention towards the former position of a barely visible Gabor patch, at various delays after its disappearance, substantially improved participants’ conscious perception of this stimulus, both on objective and subjective measures. This “retro-perception” effect was still robust 400ms after the stimulus. It has since been replicated and extended in several studies^26–29^.

Although these studies suggest that it is possible to desynchronize conscious perception from the initial sensory build-up, it might still be argued that, in these experiments, only the first feed-forward sweep of activity was triggered by the stimulus, and that retrospective cueing allowed conscious perception not by connecting sensory representations with higher-level areas, but by allowing local sensory loops to be deployed for the first time.

In the present study we push our approach much further by asking the following question: is it possible to induce retrospective conscious access to a past stimulus even after its low-level sensory representations have been degraded by masking? While such possibility is precluded by theories linking conscious perception with sensory build-up, it is a valid, albeit extreme prediction of theories advocating some independence between the two.

We therefore designed an experimental protocol which combines visual word masking with retrospective cueing by an auditory word (**Figure 1**). In visual pattern masking, conscious perception of a briefly presented written word can be drastically impaired if the word is immediately preceded and followed by other visual displays with similar low-level properties. Masking is assumed to occur because the masks replace the representation of the word within sensory areas and prevent local recurrent loops to establish a coherent representation of the word^30^; in short, masking interferes with the second processing stage. But interestingly, even when the word is rendered totally undetectable by masking, many experiments have established that it can still be processed up to a lexical and/or semantic level, presumably because the feed-forward sweep reaches high-level areas before the masks interfere with low-level representations^31,32^. Hence, after masking, what is left from the visual word is an unconscious lexical and/or semantic trace in temporal areas, while the sensory representations that led to it have been disrupted (**Figure 1**).

**Figure 1:**
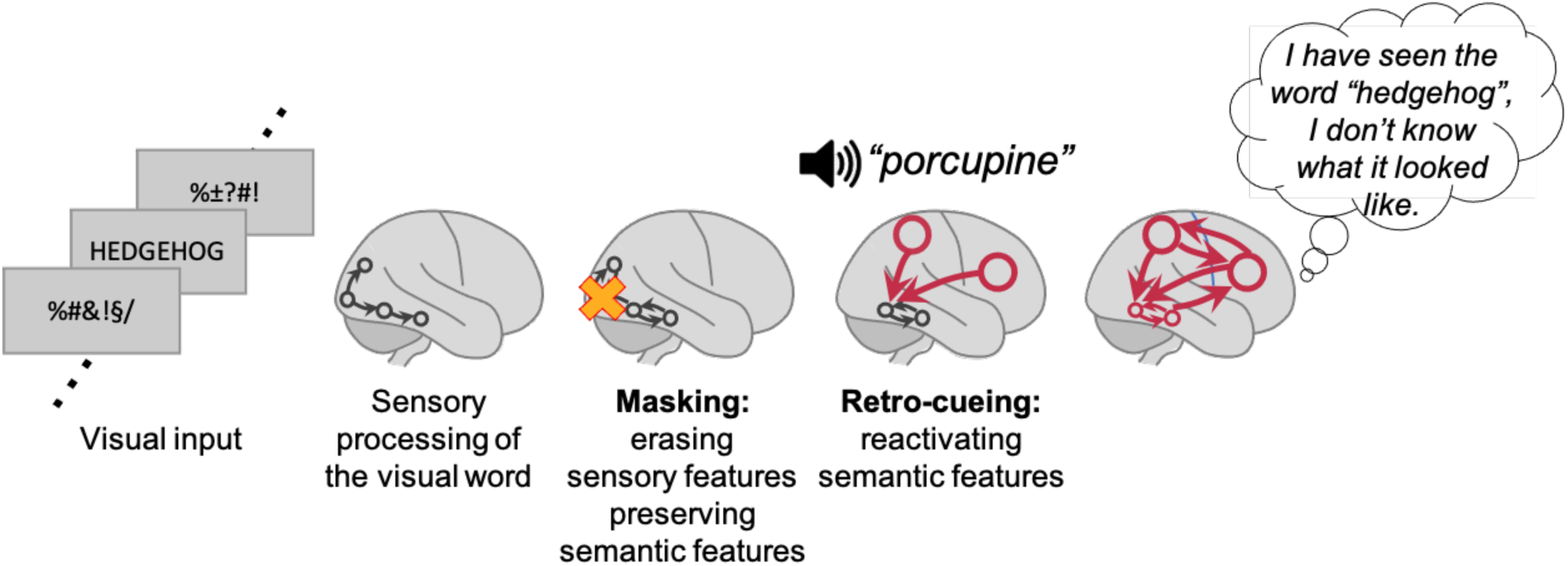
Hypothesis. This figure represents the postulated sequence of brain events associated with our protocol, combining masking and retro-cueing. When a target word is presented in a sequence of masks, it can be processed up to the level of semantic activations (first processing stage). The masks, which share many low-level visual properties with the target, act by stopping recurrent sensory processing and erasing the low-level sensory representations of the target (masking). This is what prevents conscious perception of the target word in the first place. Still, we hypothesize that a contextually congruent auditory retro-cue may bring the target word to consciousness (retro-cueing), by reactivating the latent semantic representation of the past word and hence giving it a second chance to be broadcasted through a fronto-parietal Global Workspace. This would yield conscious access to the word meaning in the absence of associated low-level information.

We know from the masked priming literature that lexical or semantic priming effects disappear for very short durations of the word, suggesting that for these too brief stimuli, unconscious processing is too weak to reach this level^33^. Hence, classical experiments showing unconscious semantic processing of masked words typically use target durations between 30 and 50ms, where they find the best compromise allowing semantic processing to occur while conscious perception is still hindered^31,33^. Here we tested a range of four different target durations (12, 24, 36 and 48ms), spanning both very short durations for which initial processing probably fails to reach semantic levels, and longer durations where unconscious semantic priming is documented.

We tested whether a subsequent semantic cue could retrospectively promote conscious access to the latent semantic representation for the durations where unconscious semantic processing occurs. The masked visual word was thus followed by an auditory word played over headphones, that could either be semantically related to it (congruent) or not (incongruent). For example, if the masked word was “hedgehog”, the retrospective cue could be either “porcupine” or “automaton” (**Figure 1**). Congruent retro-cues might induce, via semantic diffusion^34^, a top-down reactivation of the unconscious trace left by the visual word, promote the sharing of this information to a wider network of areas, and hence allow conscious access (**Figure 1**). The lexical and/or semantic representations of the masked word would thus retrospectively become conscious, but without the associated visual representations.

If such a phenomenon exists, we predict that following a congruent retro-cue, participants will be better at reporting the identity of the masked word, but will be unable to report any associated visual details, such as whether the word was written in uppercase or lowercase, or whether it has been presented above or below the fixation point. Crucially, we also predict that better identification would go hand in hand with improved conscious access, as assessed by improved subjective visibility and improved objective ability to report whether a word has been presented or not (detection sensitivity), both being classical measures of conscious access^35,36^.

We tested these predictions in a series of three experiments involving a total of 51 included participants. We then preregistered and performed two additional sets of experiments to control for a potential alternative interpretation (72 new participants). Overall, the results suggest that retrospective cues can improve conscious access to the high-level representations of a past visual word after its low-level, sensory representations have been degraded by masking. They bring support to the notion that the mechanisms of conscious access may, to a large extent, operate independently from sensory build-up.

## Results

In each experiment, the number of participants was constrained to 18 in order to ensure complete counterbalancing of stimulus presentation across participants (explained in details in **Supplementary Note 3**). This is similar to the sample size of previous experiments testing other retrospective effects which, based on a power analysis, included around twenty participants^37^. The replicability of the effects was tested across three different variants of the experiment (experiments 1 to 3), plus two sets of control experiments (experiments 4 to 7).

### Experiment 1: effect of retrospective cueing on detection, identification, and case discrimination of masked words

Due to external constraints, only fifteen volunteers could be included in Experiment 1 (see Methods). They were presented with streams of masks within which a visual word could be included, as illustrated in **Figure 2a**. The visual word could last 12, 24, 36 or 48ms, thus randomly varying stimulus strength across trials. As explained above, these values sample the steep part of the psychometric function, where unconscious semantic processing emerges. Trials where the word was absent were included to test participants’ detection sensitivity over and beyond individual response bias: indeed, comparing response distributions for present versus absent trials allows to derive the Area Under the Receiver Operating Curve for detecting the target’s presence, equivalent to a detection d’ (see Methods). Each visual sequence was followed by a retrospective cue or “retro-cue”, consisting of an auditory word that could be semantically associated to the target word (congruent trials) or not (incongruent trials) with equal probability. There was a 200ms delay between the offset of the visual word (the target) and the onset of the auditory word (the retro-cue). At the end of each trial, participants had to report first the identity of the target, by speaking into a microphone, second its case, upper or lower, using the keyboard, and third rate its overall visibility on a 9-points scale (see **Supplementary Note 1** for the full instructions). In total each participant was presented with 432 different visual words with no repetition (see **Supplementary Tables 1 and 2**). Congruency and word-pairs were balanced within and across participants as described in the Methods section. To construct our list of 432 word-pairs we performed two online free association experiments in French to complement the three existing databases. The resulting list is available in **Supplementary Table 2** and on the OSF repository for this project (https://osf.io/kd2wh/).

**Figure 2:**
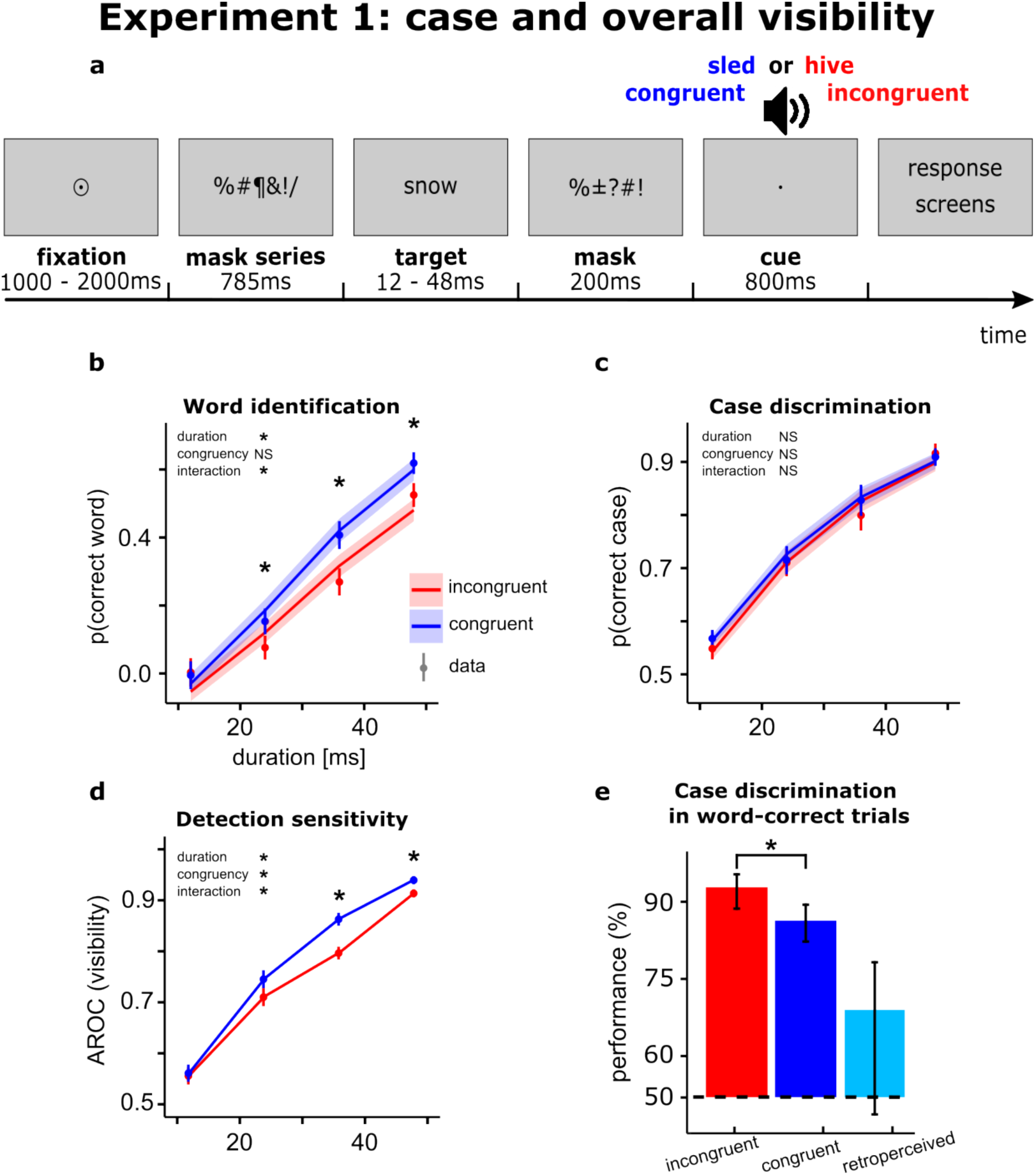
Experiment 1. **(a)** summary of trials’ structure: participants were presented with one stream of centered stimuli. Response screens asked for word identification, case discrimination and overall visibility rating, in this order. **(b)** Word identification performance, corrected for guessing (see text for details): points correspond to the data; lines represent performance estimated by the model. Error bars are standard error (SE) of the mean across observers, and error bands are the SE of model predictions across observers. **(c)** Case discrimination performance: points represent the data, lines represent performance estimated by the model, Error bars are SE of the mean across observers, and error bands are the SE of model predictions across observers. **(d)** Detection sensitivity derived from the Area under the Receiver Operating Curve of visibility ratings: error bars are standard errors of the mean across observers**. (e)** Case discrimination performance when word identification is correct, with a specific estimate for retro-perceived trials. Error bars represent 95% bootstrapped confidence intervals. In all panels, stars indicate significance of the statistical tests as described in the text.

Word identification performance and case discrimination were analyzed using a hierarchical Bayesian logistic model^38^ that included a correction for guessing for word identification (see details below). Visibility ratings and detection sensitivity derived from these ratings were analyzed using a standard frequentist approach with reported effects sizes in the form of generalized eta-squared (η²G, < 0.01 is small, > 0.14 is large^39^) for ANOVAs and Cohen’s d (*d*, < 0.2 is small, > 0.8 is large^40^) for pointwise mean differences. Wherever necessary, we used Bonferroni corrections for multiple comparisons. All tests were two-sided unless stated otherwise.

In the following, estimates from the model or their Bayesian credible intervals (BCI) are reported as odds-ratios (BCI-Odds Ratio). Note that odds-ratios represent multiplicative changes on the response: for example, an interval of [5 20] for the effect of congruency indicates that the probability of a correct response was between 5 and 20 times more likely in the congruent condition than the incongruent one. Conversely, an interval of [0.1 0.5] indicates that the probability of a correct response was 0.1 to 0.5 times more likely, in other words, 2 to 10 times less likely. Consequently, if the interval contains 1, the effect is considered not significant. Pointwise estimates of the effect of congruency at various target durations, however, are expressed as a difference in expected accuracy (BCI-diff): an interval of [0.2 0.4] for the difference between congruent and incongruent conditions indicates that the probability of a correct response was increased by 0.2 to 0.4. Consequently, if the interval contains 0, the effect is considered not significant.

#### 1.A: Word identification

The analysis of word identification performance had to include a correction for guessing. Indeed, when in doubt about the identity of the target, participants will tend to report words from the same semantic field as the retro-cue. Therefore, in congruent trials, there could be a non-negligible fraction of correct trials where participants named the target correctly just by chance. In incongruent trials, such guesses are easy to detect, because they correspond to errors. **Supplementary Figure 1** shows these guesses decrease with target duration. In order to assess whether congruency had an effect on identification performance over and beyond guessing of associated words, we first rendered the congruent and incongruent conditions comparable as far as guessing was concerned by counting as “correct” the guesses produced in incongruent trials (incorrect responses that were correctly associated with the cue) (see **Methods**). The data were fitted with mixed effect logistic regressions, with target duration, congruency and their interaction as fixed effects (see Methods). Then the mean proportion of guesses per target duration in the incongruent condition was subtracted from data points and model estimates of both conditions to obtain identification performances corrected for guessing (**Figure 2b**, see **Methods**).

Figure 2b shows the thus corrected datapoints and model’s estimates for word identification performance. We obtained a reliable fit of the psychometric functions with these logistic regressions (see diagnostics of the fits in Supplementary Table 3). Our model revealed a positive main effect of target duration (95%BCI-Odds Ratio [1.51 2.08]), no main effect of congruency (95%BCI-Odds Ratio [0.71 1.39]), and a significant interaction between target duration and congruency, with the congruency effect increasing with increasing target duration (95%BCI-Odds Ratio [1.02 1.36]). On post-hoc comparisons for each target duration, a significant effect of congruency was found for all target durations above 12ms (all the results of post-hoc statistics can be found in **Supplementary Note 2**).

#### 1.B: Case discrimination

Modeling performance for case discrimination (Figure 2c) revealed a positive effect of target duration on performance (95% BCI-Odds Ratio [1.88, 2.52]), but no effect of congruency (95%BCI-Odds Ratio [0.82, 1.50]), nor any interaction between these factors (95%BCI-Odds Ratio [0.86, 1.13]), in sharp contrast with what was observed for word identification.

#### 1.C: Detection sensitivity

We next estimated participants’ detection sensitivity (Area under the Receiver Operating Curve) (Figure 2d) by comparing the distributions of visibility ratings for target present versus target absent trials with a Mann-Whitney U-test^41^ (see Methods). A repeated measures two-way ANOVA revealed a significant increase in detection sensitivity with target duration (F(3.00, 42.00)= 92.00, p<0.001, η²G = 0.73), and congruency (F(1.00,14.00)=15.79, p<0.01, η²G = 0.04) with an interaction between these two factors (F(3.00,42.00)=3.62, p<0.05, η²G = 0.02). Bootstrapped 95% confidence intervals of the mean difference between congruent and incongruent conditions revealed a significant increase in detection sensitivity when the retro-cue is congruent for target durations of 36ms and 48ms, and a trend in the same direction for target duration of 24ms (**Supplementary Note 2**). This pattern of results suggest that congruent retro-cues improves participants’ ability at detecting the target.

**Supplementary Figure 2** provides a similar analysis on mean visibility ratings, and shows the full distributions of visibility ratings in the different conditions.

Of note, in this experiment, performances for case discrimination and detection were overall slightly higher than in the other similar experiments (experiments 3, 5 and 7), notably with performances at the shortest stimulus duration that were slightly but significantly better than chance (t(14) = 2.99 p < 0.01 for detection, and t(14) = 3.6454 p <0.01 for case discrimination). This suggests that participants in this experiment might have lower thresholds overall (the psychometric curves are displaced towards lower durations) compared to the others. This difference could be due to the fact that, for this first experiment, participants came for a pre-selection session that also constituted a 1-hour training (although on a different set of words; see Methods and Supplementary Note 4). In the subsequent studies, we improved the instructions so that participants managed to reach satisfactory levels of performance after a much shorter training and directly proceeded to the main task.

#### 1.D: Controlling for partial awareness: conditional probabilities between tasks

These results show that congruent retro-cues improve word identification, subjective visibility and detection sensitivity, but does not change performance in case discrimination. This is consistent with the hypothesis that retro-cues might induce conscious access to semantic features after the mask has degraded its low-level sensory representations, yielding a veridical increase in detection sensitivity despite no improvement in the perception of low-level features. But we should consider an alternative interpretation, in terms of partial awareness. Indeed, participants can sometimes retrieve a few letters from a masked word, even if they did not perceive it in full^42^. So, it is possible that congruent retro-cues do not act by retrospectively improving awareness of the target’s semantic attributes, but instead by allowing to combine a semantic information with existing partial awareness of some letters. For example, if participants have partial awareness of the letters “edg” and hear the retro-cue “porcupine”, they can efficiently combine both information to infer “hedgehog”.

These interpretations can be sorted out by analyzing conditional performance; if congruent retro-cues induce conscious access to the word meaning after the mask has erased information about its case, then case discrimination on these trials should be poor, and hence case discrimination for word correct trials should be worse for congruent than incongruent retro-cues. In contrast, this should not be observed if congruent retro-cues help identify the target on trials where people were already aware of a few letters.

We calculated the conditional performance at the case discrimination task when its meaning was correctly identified, separately for congruent and incongruent conditions. For this analysis, since these conditional probabilities require to examine the trial-by-trial success or failure in both tasks, we corrected for identification guesses in congruent trials on an individual trial basis, taking advantage of the trial-by-trial visibility rating: from the congruent trials where the word was correctly identified, we removed the percentage of trials with the lowest visibility corresponding to the percentage of guesses estimated from the incongruent condition (see Methods). As shown in Figure 2e, performance for case discrimination on these “word-correct” trials was very high for incongruent cues (92.9%), suggesting that, when participant reported correctly the word’s identity in this condition, they knew most of the time what was the word’s case. It significantly decreased for congruent cues, with 86.3% correct (mean difference – 7.21, 95%CI of the mean difference in %correct = [-11.66, –3.97], *d* = –1.01), confirming a dissociation of the effect of congruency on identification versus case discrimination. Finally, we can assume that, in the congruent condition, trials where the word was correctly identified include both trials where identification would have been successful irrespective of the type of retro-cue, and trials where it was successful only thanks to the congruent retro-cue (which we call “retro-perceived trials”). Using a few assumptions and rules of conditional probabilities, we could make a tentative estimate of case discrimination performance in retro-perceived trials. In Experiment 1, it was at 70.16% (mean 70.16, 95%CI = [49.01, 79.02]) (Figure 2e; see demonstration in the **Methods** and **Supplementary Figure 3**), suggesting a very degraded visual representation on those trials. Note that the width of the confidence interval shows that this estimate is noisy and only indicative.

#### 1. E: Controlling for orthographic partial awareness: orthographic distance in error trials

We conducted an additional analysis to control for a more subtle version of the partial awareness hypothesis: participants might sometimes be aware of a few letters not in their visual form, but in a more abstract orthographic form. For example, participants might be aware that the triplet of letters “e, d and g” was presented without having seen their visual attributes; this information might nevertheless combine with the congruent semantic retro-cue to produce successful identification.

We tested for this possibility by examining the orthographic distance between participants’ response and the actual target in error trials, using the Levenshtein distance which estimated the number of substitutions, additions and deletions required to match the participant response to the correct answer (**see Methods**). Indeed, if orthographic partial awareness exists in our protocol, then participants’ responses should include these orthographic fragments even when they fail to correctly identify the word. For example, if they have perceived “edg” they might report “knowledge” instead of “hedgehog”. Furthermore, if improved word identification with congruent retro-cues is due to these trials, then their proportion among error trials should decrease for congruent compared to incongruent retro-cues.

As shown in **Supplementary Figure 4**, orthographic distance in error trials decreased significantly with increasing target duration (F(3,42)=40.16, p< 0.001, η²G = 0.42), confirming that there was some partial orthographic awareness in this protocol, and that it increased with increasing target duration. However, there was no significant effect of congruency on this orthographic distance (F(1,14)=0.05, p=0.82) and no significant interaction between congruency and target duration (F(3,42)=1.06, p=0.37), suggesting that the congruency effect we observed on identification did not rely on partial orthographic awareness.

#### 1. F: Conclusion

In conclusion, we observed that congruent retro-cues improved participant’s identification as well as their sensitivity to detect the presence of the words (Figure 2b and d), without improving their ability to report their case (Figure 2c). This suggests that congruent retro-cues might induce conscious detection and conscious identification of stimuli whose low-level features can no longer be reported because of the masks. We made sure that this effect could not be accounted for by partial awareness. Next, we conducted two additional experiments with small variations from this protocol in order to test the replicability of these effects and their generality. We also took advantage of these experiments to further characterize the associated subjective experience. In Experiment 2 we changed the visual task from case to position discrimination, and in Experiment 3 we collected ratings about the confidence in the response instead of the visibility of the word.

### Experiment 2: effect of retrospective cueing on detection, identification, and position discrimination of masked words

In Experiment 2 we tested whether performance dissociation between semantic and physical properties extended to another fundamental visual attribute: the position of the visual word on screen. Participants were presented with two streams of masks appearing above and below the central fixation point (Figure 3a). On each trial, the target word could appear in either stream, and participants had to report the target’s position as well as its identity.

**Figure 3:**
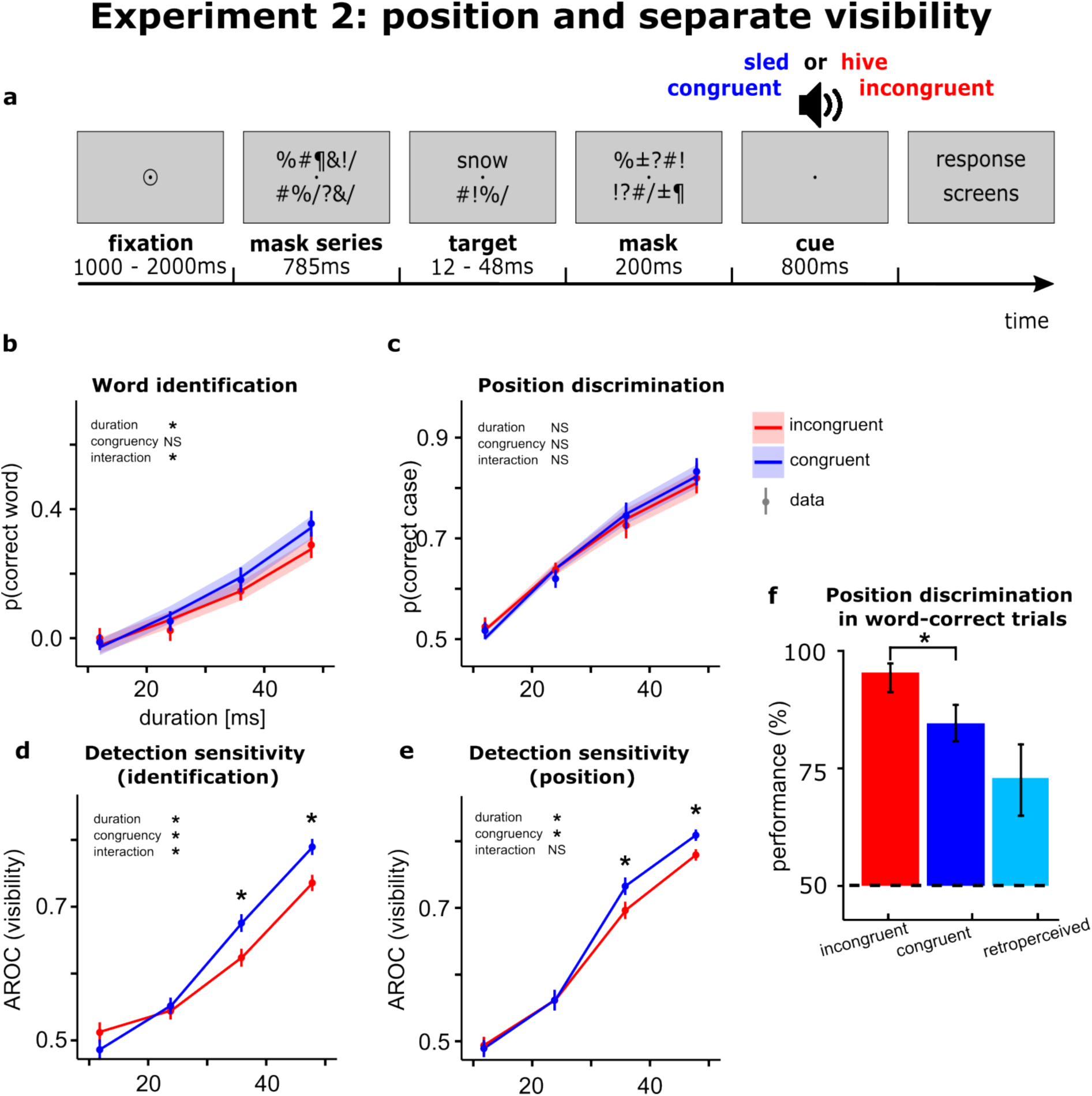
Experiment 2. **(a)** summary of the trials structure: participants were presented with two streams of stimuli above and below fixation. Response screens asked for position discrimination, visibility on position, word identification, and visibility on word identity, in this order. **(b)** Word identification performance corrected for guessing. **(c)** Position discrimination performance. **(d)** Detection sensitivity derived from visibility ratings of word identity. **(e)** Detection sensitivity derived from visibility ratings of word position. **(f)** Conditional position discrimination performance in trials with correct word identification. All conventions as in Figure 2.

We also explored in more details how retro-cueing affected subjective visibility by asking participants to provide visibility ratings along both high-level and low-level features. At the end of each trial participants first reported the position of the target word (above or below fixation), and then provided a visibility rating of the word’s position on a scale from 0 to 8; then they reported the target’s identity by speaking into a microphone, and again provided a visibility rating of the word’s identity on a scale from 0 to 8. The instructions for the two visibility scales put different emphasis on the position versus identity of the word, but both requested that the participant report the sensation of having seen a word versus not at all (distinction between zero visibility and non-zero visibility, **Supplementary Note 1**). This exploratory manipulation aimed at testing whether participants could distinguish over which dimension their subjective visibility increased. If they could, the prediction the effect of congruency on subjective visibility and derived detection sensitivity should only be observed for word identification, not position. However, we were also interested in another possibility, that participants might have had an illusion of seeing the low-level feature when they felt they detected the stimulus, in which case this dissociation in subjective visibility along the two dimensions should not be present.

Eighteen participants took part in this experiment. When analyzing the results, we first made sure that participants were not biased into attending a specific stream by analyzing their position response on target-absent trials. Participants absolute deviation from an ideal 50% of each type of response in catch trials was 14.58±9.20% (95% CI [10.76 18.98], maximum deviation 31.25%), presumably reflecting individual biases towards one stream or the other in the absence of an overt strategy.

#### 2. A: Word identification

The main analyses were conducted as in Experiment 1 (see **Methods**): for word identification (Figure 3b), we evidenced a positive effect of target duration (95%BCI-Odds Ratio [1.34 1.80]), and a significant interaction between congruency and target duration (95%BCI-Odds Ratio [1.002 1.27]) but no main effect of congruency (95%BCI-Odds Ratio [0.63 1.19]). None of the post-hoc pointwise comparisons at each target duration reached significance (the results of all post-hoc comparisons can be found in **Supplementary Note 2**). This pattern of results suggests that the interaction effect on word identification was present in Experiment 2 but was less strong than in Experiment 1.

#### 2.B: Position discrimination

For position discrimination (Figure 3c), performance increased significantly with longer target durations (95%BCI-Odds Ratio [1.78 2.24]), but there was no effect of congruency (95%BCI-Odds Ratio [0.91 1.56]) and no interaction (95%BCI-Odds Ratio [0.87 1.09]), suggesting that, in contrast to word identification, congruency had no effect on position discrimination.

#### 2.C: Detection sensitivity

Detection sensitivity derived from the first visibility scale, related to word identification (Figure 3d), showed a significant effect of target duration (F(3.00,51.00)=38.84, p<0.001, η²G=0.48), congruency (F(1.00,17.00)=9.24, p<0.01, η²G=0.01), and a significant interaction (F(3.00,51.00)=8.50, p<0.001, η²G=0.02). Post-hoc pointwise comparisons at each target duration showed a significant increase in performance with congruency at durations 36ms and 48ms (**Supplementary Note 2**). Detection sensitivity analysis for the other visibility scale, related to word position (Figure 3e), showed a significant effect of target duration (F(3.00,51.00)=48.61, p<0.001, η²G=0.54), a significant yet very small effect of congruency (F(1.00,17.00)=7.53, p=0.01, η²G=0.005), and no interaction (F(3.00,51.00)=2.35, p=0.08). Pointwise comparisons at each target duration showed an increase in detection sensitivity with congruency at durations 36ms and 48ms (**Supplementary Note 2**). Finally, we conducted a 2-way repeated measures ANOVA on mean difference in detection sensitivity between congruent and incongruent conditions, by target duration and type of visibility rating (related to word identification or position discrimination) to assess whether congruency affected detection sensitivity differently according to type of stimulus dimension that was emphasized. This analysis did not reveal a main effect of type of task (F(1.00,17.00)=0.52, p=0.48), nor an interaction between task and target duration (F(3,51)=1.72, p=0.17), suggesting that both visibility ratings were not differently affected by congruency across experimental conditions.

This pattern of result is somewhat intermediate between the two initial predictions we could make: there was no clear dissociation in the use of the visibility scale for identification and position, but still there was a slight trend towards a weaker effect of congruency on position (Figure 3d-e). This could suggest that improved perception of the word identity with congruent retro-cues is accompanied by an illusion of having seen its position better despite no objective improvement on this dimension. Another possibility, however, is that the instructions for the two scales were not sufficiently distinct: indeed, both scales shared the instruction to use visibilities bigger than zero as soon as they had a feeling that they might have detected something.

**Supplementary Figure 5** provides a similar analysis on mean visibility ratings, and shows the full distributions of visibility ratings in the different conditions.

#### 2.D: Controlling for partial awareness: conditional probabilities between tasks

As for Experiment 1, we next addressed possible alternative interpretations involving partial awareness. If word identification improvement was due to partial awareness of a few letters with their physical attributes, then position discrimination for trials where the word was correctly reported should be at least as good on congruent trials than on incongruent trials. We corrected for guesses on a trial-by-trial basis as in Experiment 1. Position discrimination performance in word-correct trials (Figure 3f) decreased from 98.5% in incongruent trials to 85.7% in congruent trials (mean difference congruent minus incongruent: –11.88%, 95%CI [-15.87 –8.45], *d* = –1.60), arguing against an interpretation in terms of partial awareness of physical attributes. Tentative estimate of position discrimination performance in retro-perceived trials (same methods as in Experiment 1) was 74.89%, significantly above chance (95%CI [66.82 82.05], *d* = 4.39), denoting the fact that the perception of position was very degraded, although not null, on retro-perceived trials. Note that the width of the confidence interval shows that this estimate is noisy and is only indicative.

#### 2.E: Controlling for orthographic partial awareness: orthographic distance in error trials

We next tested if partial orthographic awareness of the target word could account for our results. As explained in Experiment 1, under this hypothesis the proportion of errors that share orthographic constituents with the masked word should decrease for congruent compared to incongruent retro-cues, hence increasing the orthographic distance between the erroneous responses and the true responses. As shown in **Supplementary Figure 4**, no such effect was observed: the repeated measures ANOVA (duration x congruency) showed an effect of duration (F(3,51)=3.584, p=0.02, η²G = 0.05) but no effect of congruency (F(1,17)=0.209, p=0.65) nor an interaction (F(2.23,37.93)=1.150, p=0.34), suggesting that this partial awareness interpretation cannot account for the results.

#### 2.F: Conclusion

In conclusion, Experiment 2 replicates the effect of retrospective cueing on word identification and word detection observed in the previous experiment, and successfully extends the dissociation between awareness of abstract features versus physical features to the spatial location of the stimulus. However, overall performance as well as the effect of congruency was reduced at the word-identification task, possibly due to participants attending two locations on screen compared to one in Experiment 1. Furthermore, the results obtained when distinguishing visibility ratings for identification and for position could have different interpretations, an issue we address in Experiment 3.

### Experiment 3: effect of retrospective cueing on identification, case discrimination and confidence in both tasks

We designed Experiment 3 to provide an additional replication of our initial results (Experiment 1) and probe a possible distinction between subjective visibility ratings and confidence ratings. Indeed, in Experiment 2, detection sensitivity, as measured with visibility ratings, increased similarly with congruent retro-cues for both visual (position) and more abstract (semantic) dimensions of the stimulus, contrasting with the dissociated effects of congruency on word identification versus position discrimination. This could have two interpretations: either improved perception of the word identity with retro-cues is accompanied by an illusion of better perceiving its visual features, or the partially shared instructions for both visibility ratings induced this similarity in responses. Here we address this issue by changing visibility scales for confidence scales, with each scale being dedicated to judging confidence over either the identification response or the case response. If the similar patterns obtained for visibility ratings was due to overlapping instructions, these well-separated confidence ratings should instead reflect the dissociation observed in first-order performance.

Eighteen new participants took part in this experiment. The structure of the experiment was the same as in Experiment 1, as illustrated in Figure 4a, except that, at the end of each trial participants had to report the case of the target word, followed by their confidence on this response, and then they had to report the word identity in a microphone, followed by their confidence in this response. On both confidence scales “0” corresponded to an absence of confidence in the corresponding response, regardless of whether they perceived the stimulus or not (**Supplementary Note 1**).

**Figure 4:**
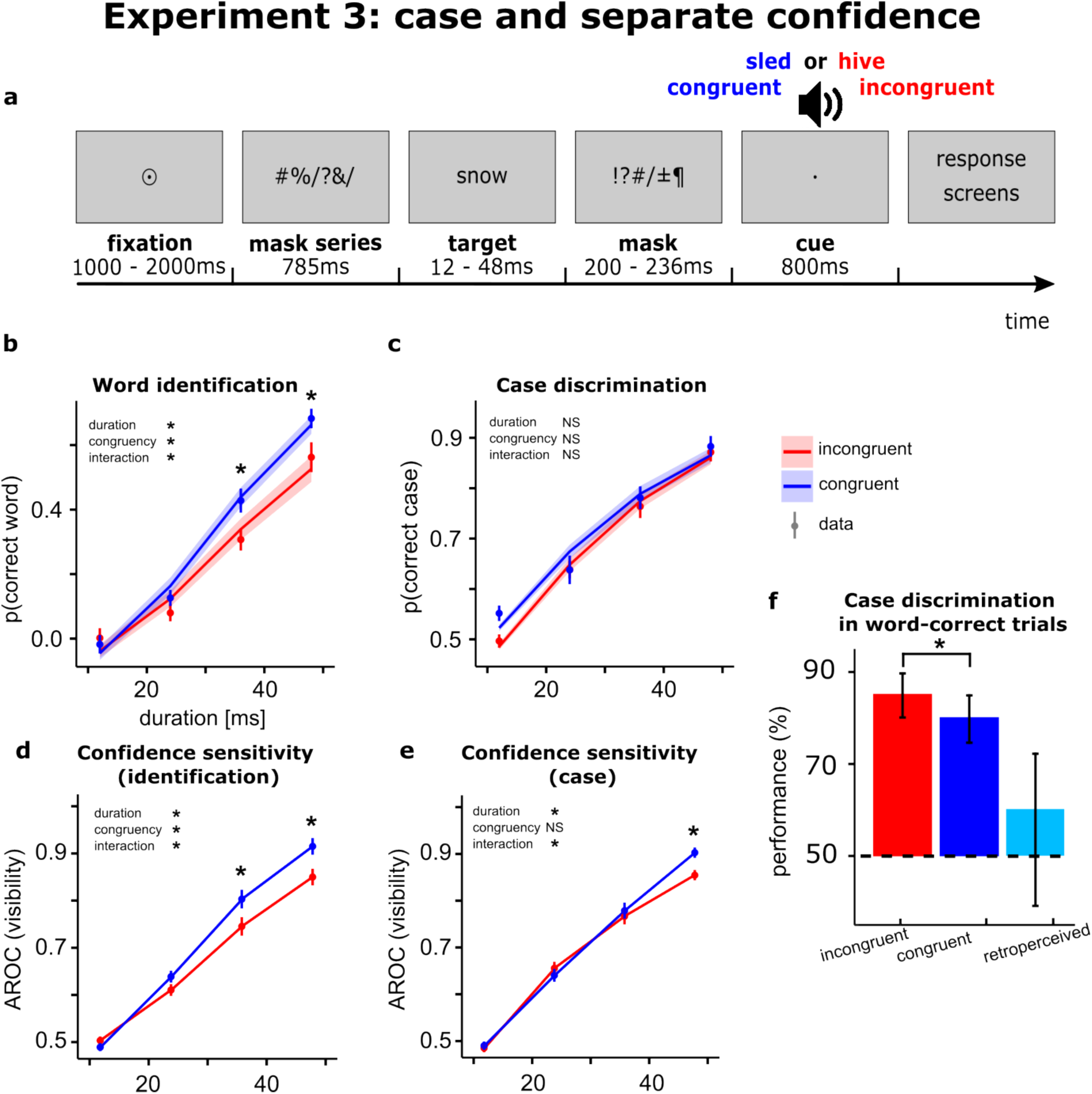
Experiment 3. **(a)** summary of trials’ structure: participants were presented with one stream of centered stimuli. Response screens asked for: case discrimination, confidence on case, word identification, and confidence on word identity, in this order. **(b)** Word identification performance, corrected for guessing. **(c)** Case discrimination performance. **(d)** Confidence sensitivity for words, derived from the AROC of confidence ratings over the word identification response. **(e)** Confidence sensitivity for case, derived from the AROC of confidence ratings over the case discrimination response. **(f)** Response consistency probability. **(g)** Conditional case discrimination performance. All errors bars as in Figure 2.

#### 3.A: Word identification

The main analyses were conducted as in Experiment 1: for word discrimination (Figure 4b), there was a positive effect of target duration (95%BCI-Odds Ratio [1.75 2.65]), congruency (95%BCI-Odds Ratio [1.02 1.50]), as well as a significant interaction between these two factors (95%BCI-Odds Ratio [1.16 1.50]). Post-hoc pointwise comparison at each target duration showed an increase in performance with congruency at durations above 24ms (**Supplementary Note 2**).

#### 3.B: Case discrimination

For case discrimination (Figure 4c), we found an effect of target duration (95%BCI-Odds Ratio [1.45 1.92]), but no effect of congruency (95%BCI-Odds Ratio [0.69 1.16]) and no interaction (95%BCI-Odds Ratio [0.95 1.18]). This pattern of results replicates the dissociation observed between word identification and case discrimination in Experiments 1 & 2.

#### 3.C: Confidence sensitivity

Confidence sensitivity analysis for word identification (Figure 4d**, see Methods**) showed a significant effect of target duration (F(1.03,17.64)=121.19, p<0.001, η²G = 0.73), congruency (F(1.00,17.00)=14.90, p<0.01, η²G = 0.04), and a significant interaction (F(3.00,51.00)=6.26, p<0.001, η²G = 0.03). Pointwise comparisons at each target duration showed an increase with congruency at durations 36ms (**Supplementary Note 2**) and 48ms. This shows that confidence sensitivity to word identification reflects the boost to word identification provided by congruent cues.

Confidence sensitivity analysis for the case discrimination response (Figure 4e) showed an effect of target duration (F(3.00,51.00)= 96.94, p<0.001, η²G = 0.67) and an interaction (F(2.17,36.98)=4.45, p<0.001, η²G = 0.01), but no main effect of congruency (F(1.00,17.00)=2.84, p=0.11). Pointwise comparisons at each target duration showed an increase in case confidence sensitivity with congruency at duration 48ms (**Supplementary Note 2**). This suggests a visibly smaller effect of congruency on confidence sensitivity in this case, namely that participants slightly increase their confidence on case discrimination for congruent trials at the longest target duration, despite an absence of effect of congruency on first-level performance.

Finally, we conducted a 2-way repeated measures ANOVA on mean difference in confidence sensitivity between congruent and incongruent conditions, by target duration and type of visibility rating (related to word identification or position discrimination) to assess whether congruency affected confidence sensitivity differently according to the dimension that was emphasized. This analysis revealed a main effect of type of confidence scale (F(1.00,17.00)=8.30, p=0.01, η²G = 0.03), as well as an interaction between type of confidence scale and target duration (F(3,51)=7.79, p<0.01, η²G = 0.48) on this congruency effect. This contrasts with the absence of significant interaction between congruency and type of scale on detection sensitivity in Experiment 2.

**Supplementary Figure 6** provides a similar analysis on mean confidence ratings, and shows the full distributions of visibility ratings in the different conditions. A closer look at these distributions shows that the slight effect observed at 48ms for confidence sensitivity relates to an increase in confidence for ratings that were already confident (from confidence 6 to confidence 8), not a change from low confidence to high confidence, in stark contrast with the distributions observed for word confidence.

Overall, these results show that when participants were asked to rate their confidence separately on each dimension their ratings reflected the differential effect of congruency on identification versus case discrimination, suggesting that, although congruent retro-cues made them detect the stimulus presence better, this was not accompanied by an illusion of confidence about their ability to report the word’s case. The similar patterns observed over the two dimensions when using visibility ratings in Experiment 2 was thus probably due to a partial overlap in the instructions for reporting visibility over the two dimensions.

#### 3.D: Controlling for partial awareness: conditional probability analysis

Case discrimination performance in word-correct trials – corrected for guessing – (Figure 4f) was significantly decreased in congruent trials compared to incongruent trials (mean –6.27%, 95%CI [-9.53 –3.32], *d* = –0.58). Tentative estimate of case discrimination performance in retro-perceived trials was not significantly different from chance (mean 61.93%, 95%CI [45.76, 72.53]). In accordance with what was observed in the previous experiments, this suggests that the additional identifications brought about by congruent cues might rely on trials where information on the word’s case is strongly degraded or absent. Note that the width of the confidence interval shows that the estimate of performance for retro-perceived trials is noisy and is only indicative.

#### 3.E: Controlling for orthographic partial awareness: orthographic distance in error trials

As in previous experiments, we observed a decrease in orthographic Levenshtein distance in error trials with target duration (F(2.17,36.93)=22.790, p<0.001, η²G = 0.25), but no effect of congruency (F(1.00,17.00)=1.891, p=0.19, η²G = 0.08) nor an interaction (F(1.68,28.57)=0.960, p=0.39, η²G = 0.01) (**Supplementary Figure 4**), contrary to what would be expected if the effect of congruent retro-cues relied on orthographic partial awareness.

#### 3. F: Conclusion

In conclusion this experiment replicated our initial results and suggested again that congruent retro-cues induce a form of dissociated conscious access to the word’s semantic representation in the absence of sensory representation. Furthermore, while previous experiments suggested that this was accompanied by increased subjective visibility and increased detection sensitivity, the present results indicate that this is not conflated with a strong illusion of confidence about the ability to report the word’s case. We next performed 4 preregistered control experiments to further assess whether this performance dissociation was sensitive to the temporal order between cue and target.

### Experiments 4 through 7: comparing the effect of pre-cues and retro-cues

In these last experiments, we tested whether performance dissociation between high-level and low-level information was specific to retrospective cueing. Our main hypothesis would predict so, since we postulate that this dissociation arises because the retro-cue triggers conscious access after low-level features have been disrupted by masking. However, as mentioned earlier, an alternative interpretation of our results could be that the cue does not change the awareness status of the various representations of the stimulus, but only allows better identification and detection performance by combining partial information on the word with a semantic context. If this were the case, then the temporal order of the visual and auditory presentation should not matter: in both cases the combination of information should occur, producing the same dissociation. Alternatively, if the retro-perception interpretation is correct, we predict that the dissociation should be unique to retro-cueing, while pre-cueing should alleviate masking at both the low and high level.

Four experiments were specifically designed and preregistered to investigate this question by comparing the effect of pre-cues (Experiment 4 and 6) and retro-cues (Experiment 5 and 7) (https://osf.io/cnf68/ and https://osf.io/8rfw7/). Eighteen participants were included in each of these experiments (total 72, see Methods for details). We took advantage of these experiments to control for additional factors from our initial design. First, to ensure that the observed improvements in performance were due to a positive effect of congruent cues and not, or not only, due to a negative effect of incongruent ones (priming the incorrect lexicon could be thought to decrease performance), we replaced incongruent cues by neutral non-words that were either the reversed audio track of congruent cues (4 and 5) or pure tones with the same duration and envelope as the congruent cue (6 and 7). We also changed cueing from associative (“porcupine” – hedgehog) to repetition (“hedgehog” – hedgehog) in all Experiments. Consequently, since word identification became trivial when the cue actually repeated the target, participants only had to report letter-casing and visibility of the whole stimulus. This maximized our chance to observe cueing effects at the level of visual representations, since participants were focused on the visual attributes of the target. Neutral cues were changed between Experiments 4&5 and 6&7 because participants sometimes reported being able to recover the original word from the reversed audio tracks on neutral trials, hence bringing neutral trials closer to congruent trials. The patterns of results were similar for both pairs of experiments (see **Supplementary Figure 7**). In what follows the two pre-cueing experiments (4 and 6) and the two retro-cueing experiments (5 and 7) were pooled for analysis.

Detection sensitivity (derived from visibility ratings) was analyzed similarly to previous experiments. Case discrimination was analyzed using separate logistic regressions for pre-cueing and retro-cueing experiments, using target duration and congruency as factors. These analyses were followed by a third logistic regression model that included the type of cue (pre or retro-cue) as a factor, alongside congruency and target duration. One-sided tests were used, given that our previous experiments allowed strong priors about the directions of the expected effects (positive effect of congruent cues vs incongruent ones, positive effect of increasing target duration, negative effect of retro-cueing vs pre-cueing). Results of both experiments are presented in Figure 5.

**Figure 5:**
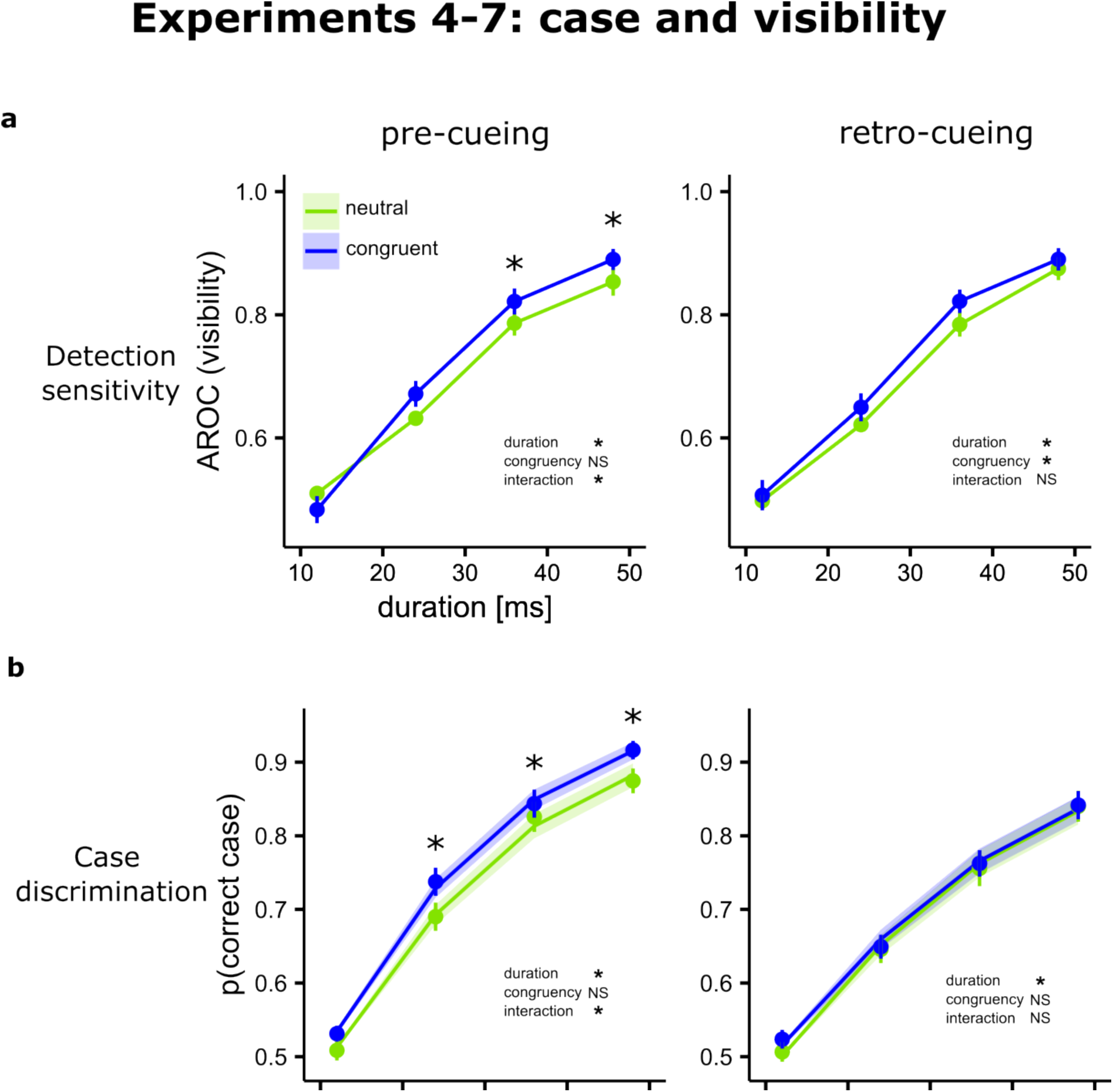
Experiments 4 – 7. Detection sensitivity **(a)** and case discrimination performance **(b)** are shown for pre-cueing (left panels) and retro-cueing (right panels). Stars indicate intervals where post-hoc comparisons were significant or panels where the statistical analysis showed a significant effect of congruency with no significant post-hoc test, vertical lines are standard error of the mean across participants (lower panels), or the mean predictions of the model (lower panels).

For detection sensitivity, the patterns obtained matched our predictions as both pre and retro-cueing influenced detection (Figure 5a). For pre-cueing, detection sensitivity significantly increased with target duration (F(1.89,66.00)= 236.59, p<0.001, η²G = 0.62). There was no main effect of congruency with a marginal p-value (F(1,35)=3.34, p=0.08) and a significant interaction between duration and congruency (F(3,105)=9.37), p<0.001, η²G = 0.01). Further one-sided post-hoc pairwise comparisons showed an increase with congruent pre-cues for target durations 36 and 48ms (**Supplementary Note 2**). For retro-cueing, we found a significant main effect of target duration on detection sensitivity (F(1.54,54.06)=202.96, p<0.001, η²G = 0.64), a significant main effect of congruency (F(1,35)=5.83, p=0.02, η²G = 0.01), and no significant interaction between duration and congruency (F(1.54,54.06)=1.70, p=0.1). Hence, although one was reflected in an interaction and the other in a main effect, in both cases congruent cues significantly improved detection sensitivity.

For case discrimination (Figure 5b), the pattern of results also matched our predictions: in analyses conducted separately for the two types of cues, we found that congruency improved case discrimination for pre-cues as evidenced by a significant interaction with target duration (target duration 95%BCI-Odds Ratio [1.99 2.42], congruency 95%BCI-Odds Ratio [0.80 1.14], interaction 95%BCI-Odds Ratio [1.04 1.23]), but for retro-cues we found no significant effect of congruency and no interaction with duration (target duration 95%BCI-Odds Ratio [1.73 2.13], congruency 95%BCI-Odds Ratio [0.94 1.30], interaction 95%BCI-Odds Ratio [0.90 1.04]). Data were then analyzed using the full model (congruency x target duration x type of cue) which revealed a main effect of target duration (95%BCI-Odds Ratio [2.00 2.47]), an interaction between congruency and target duration (95%BCI-Odds Ratio [1.02 1.19]), target duration and type of cue (95%BCI-Odds Ratio [0.74 0.99]), and, crucially, a triple interaction between target duration, congruency and type of cue (95%BCI-Odds Ratio [1.02 1.20]). No main effect of congruency (95%BCI-Odds Ratio [0.84 1.19]) or type of cue (95%BCI-Odds Ratio [0.91 1.35]), or interaction between congruency and type of cue (95%BCI-Odds Ratio [0.86 1.39]) were found.

Overall, this pattern of results suggests that, while both congruent pre-cues and retro-cues improved detection, only pre-cues, not retro-cues, improved case discrimination. This validated our predictions, confirming that the dissociation between detection and case discrimination is specific to retro-cueing. Of note, the effect of retro-cues on detection sensitivity was smaller than in Experiments 1-3. This could be explained by the fact that, in the present control experiment, participants were not performing a task on the cued dimension of the stimulus (its meaning), as was the case in previous experiments, or to the fact that congruent and incongruent cues could be distinguished. It could also suggest that, in previous experiments, incongruent cues could have had a small hindering effect compared to a neutral condition.

## Discussion

In this study we probed a counter-intuitive prediction, derived from the global neuronal workspace theory, that it should be possible to trigger conscious access to high-level representations of a stimulus even after the associated low-level sensory representations have been masked. We conducted a first series of three experiments which all replicated the same observations: over a range of several durations of a masked visual word, on the rising part of the psychometric function, participants’ ability to report the identity of the word was substantially improved when it was followed by a semantically related auditory word (congruent retro-cue) compared to an unrelated word (incongruent retro-cue). The range of durations where this improvement occurred matched the typical durations where unconscious semantic processing is known to take place^31,33^. This improvement was accompanied by increased objective ability to detect the presence of the target, and increased subjective visibility, showing that the effect did not rely on subliminal processing but rather on the fact that retro-cues retrospectively improved conscious access. In contrast, no improvement was observed for reporting lower-level features such as letter casing (Experiments 1 and 3) or position on screen (Experiment 2). Further analyses confirmed that congruent retro-cues induced a dissociation between the ability to identify the word and to report its visual features. Experiments 5 and 7 replicated the observation that retro-cues induced increased detection sensitivity and visibility, but had no effect on case discrimination.

Although these results verified our initial predictions, we had to consider another possible interpretation. Congruent retro-cues could have improved performance not by retrospectively improving awareness but by providing additional information on trials where participants already had some partial awareness of the masked word^42–44^. For example, an initial glimpse of the letters “edg”, combined with the retro-cue “porcupine”, could allow to infer “hedgehog”. We controlled for this at three different levels. First, if better identification occurs on trials where people saw a few letters, congruent retro-cues should increase the association between the ability to report the word’s visual features and its identity. But here we observed the opposite effect (Figures 2e**, 3f, 4f**). Second, we controlled for a more subtle version where partial awareness would concern abstract, non-visual, orthographic features. Contrary to what would be predicted in this case, retro-cueing did not influence the orthographic similarity between the target and the words produced by participants on error trials (**Supplementary Figure 4**). Finally, we preregistered and performed two series of two experiments each to test the importance of temporal order between masking and cueing. Under our initial hypothesis, performance dissociation should critically depend on order, while order should not matter under the partial awareness hypothesis, which assumes that better performance only arises from combining the cue with partial information about the target. The results validated our predictions that pre and retro cues impact performance differently (Figure 5): while pre-cues improved both detection and case discrimination, retro-cues deployed on the same protocol improved participants’ detection without improving case discrimination, replicating again our previous observations. These controls hence emphasize that the dissociation between detection and case discrimination specifically arises when the cue is placed after the target and its masks, validating our proposition that the impact of retro-cues on conscious detection occurs after low-level features of the target have been degraded by masking.

Several previous studies have shown that a retrospective context word can influence the lexical processing or identification of a preceding masked word^45–50^. Here we further investigated its impact on conscious access by using two benchmark measures^51^: objective detection sensitivity (or d’) and subjective visibility. Our results showed that improved identification with retro-cueing was not akin to a form of “blindsight”^52,53^, but that it was instead genuinely associated with a change in participants’ conscious experience, as reflected by increased subjective visibility and objective detection sensitivity.

We further attempted to probe whether participants were able to distinguish over which dimension, high-level or low-level, their subjective visibility increased. We obtained mixed results: in Experiment 2, an effect of retro-cue congruency could be observed on subjective visibility and derived detection sensitivity for both dimensions, although the effect was a bit weaker for the low-level dimension (Figure 3d**,e**). One interpretation could be that improved detection of the word meaning was accompanied by an illusion of having seen also lower features of the stimulus despite no effect on objective performance; however, this result could also be due to partially shared instructions since, for both scales, participants were instructed to use visibilities bigger than zero as soon as they had a feeling that they might have detected something. In Experiment 3 we hence replaced visibility for confidence and, this time, each scale was clearly associated with judging confidence over either the identification response or the case response. We observed a dissociation in confidence, matching the dissociation observed on first-level performance (Figure 4d**,e**). There was still a slight effect of congruency for the longest duration for confidence in the low-level response: on the confidence distributions, this corresponded to a transfer of trials with high confidence to trials with maximal confidence, in contrast with the effect observed for word identification, where we observe a transfer of responses from low and mid-level confidence to maximal confidence **(Supplementary Figure 6**). Overall, this suggests that participants are generally willing to express better perception of the stimulus when using a visibility scale, especially given partially shared instructions across the different scales, but that they probably do not experience an illusion of seeing the low-level feature, as the effect of congruency on this dimension only concerns a modulation of confidence on trials that were already confident.

However, future work will be needed to better characterize participants’ subjective experience in the protocol presented here. As highlighted in studies on sound induced flash illusions, metacognitive awareness can be much more subtle and difficult to reveal than first order effects on behavior, especially in rapid multisensory integration situations^54,55^. In our case, null-effects for subjective visibility could be specifically reassessed in a dedicated protocol: we provided instructions with a focus on distinguishing between the presence and absence of the stimulus, which may not have been suited to explore subtle differences in the perceptual quality of experiences of word-meaning and word-form.

These results bring important novel evidence for current debates about the role of sensory processing in conscious perception. They seem to challenge theories postulating that conscious perception is linked with the establishment of local loops within sensory cortices^10,12^, since visual masking typically disrupts these loops, and hence should definitively preclude conscious perception thereafter^30,33,56^. Instead, they appear to validate a radical prediction of contender theories, and corroborate the neurophysiological mechanism proposed in Figure 1. We know from previous literature that the processing of a visual word can reach lexical or semantic analysis before masking disrupts the associated low-level representations^33,57,58^. Although these high-level representations can prime subsequent conscious words, they typically remain unconscious. Here we propose that retrospective cueing triggers conscious access to these initially unconscious high-level representations via the following mechanism: first, the retro-cue induces a spreading of activations in lexical and/or semantic networks^34^; if the unconscious high-level representations left by the past visual word belong to the same lexical field, this should reactivate them^59–62^. These reactivated representations being relevant to the task, they might be selected by attention and broadcast within a global workspace^9,17^, allowing for conscious access despite the degradation of the associated visual details, several hundred milliseconds after stimulus disappearance.

Although these results seem to contradict classical formulations of local loop theories which typically focus on the construction of sensory representations within the first levels of the hierarchy –that are disrupted by pattern masking^4,56,63^, we could still interrogate whether an extended formulation of this local loops framework could, in hindsight, accommodate our results. One could argue that retro-cues act by retrospectively triggering local recurrent processing in higher-level areas such as temporal visual areas. We would need to assume that, although our masks only disrupt recurrent processing at low levels, they nevertheless manage to affect recurrency also at higher levels, and that retro-cues allow to establish these initially failed local recurrencies. The role of local recurrencies in the build-up of sensory representations is well established for low-level sensory attributes (typically a figure-ground context modulates the encoding of primary visual attributes via these recurrencies)^4,56,63^, and it is this type of phenomena that has grounded the local loops theory. It is less clear what type of additional local recurrencies would be required at the lexical or semantic level for allowing conscious perception of representations at this level only, knowing that unconscious lexical or semantic representations of the masked word exist in the first place, independently of retro-cueing. So, while it remains possible to interpret these results within an extended formulation of the local loops framework, such interpretation would be less straightforward.

Importantly, over and beyond arbitrating between different models, these results bring about important novel results regarding the flexibility of conscious access mechanisms with regards to the initial stages of sensory build up that any model of conscious perception should take into account. They corroborate the previously described retro-perception phenomenon, namely the possibility to trigger conscious perception retrospectively^26,28,29,37,64,65^, and extends it to visual masking. In doing so it shows that conscious access is not only flexible in time but also in type of content. Indeed, present results reveal a novel phenomenon of dissociated conscious percepts, where participants report having seen a stimulus that they correctly identify, despite being almost clueless about its visual details. This suggests that the mechanisms for conscious access are relatively insensitive to the “completeness” of an experience, so that they can operate on higher-order representations in the absence of a precise sensory base. This is reminiscent of the large inter-individual variations in how a representation can be experienced, for example in aphantasia, an absence or near-absence of mental imagery which concerns about 2.5% of the population^66^. These kinds of phenomena are all the more interesting in that they challenge our intuitions about conscious perception and its relationship with conscious experience in general.

These results also resonate with the Reverse Hierarchy Theory^67^, which proposes that implicit feedforward processes trigger the appropriate receptive fields in higher-level areas for fast and effortless gist-perception (here, word meaning), and that conscious perception of details (here, letter casing or location) may be optional and requires focused reactivation of low-level areas from the top. In Reverse Hierarchy terms, the present results would suggest that retro-cues induce top-of-the-hierarchy conscious experiences followed by a downward stream of detail enhancement, which fails when it reaches levels affected by masking.

Along a similar line, these results also extend the idea of a “predictive brain”^68^. There is now ample evidence that what we will perceive next, and even *whether* we will perceive it consciously or not^69^ is strongly influenced by the past, and by a projection of what can be expected next based on that past. For example, Lupyan & Ward show that semantic prediction influences whether a stimulus crosses the consciousness threshold or not^69^. Here we show that combining information to get more than instantaneous evidence is a process that can go in both directions: the predictive brain is also a retrodictive brain.

In conclusion, these findings are highly relevant to current debates over the role of sensory processing in consciousness. They demonstrate the extreme flexibility of the form that can be taken by conscious perception and conscious access to an external stimulus, both in terms of timing and content. They suggest that conscious perception of a word can be reduced to its most essential property, its meaning: we know that a stimulus has been presented; we know what it is; we agree to say that we have “seen it”, but we don’t know what it looks like. This brings support to theories of consciousness for which the processes of becoming conscious of a representation can be distinguished from the processes underlying its elaboration, suggesting a wide range of ways to experience any single event. More generally, these results challenge our understanding of the relationship between conscious experience and external events, and interrogate the very nature of what we call a conscious percept. They also open new perspectives and novel protocols for exploring the neural underpinnings of conscious processing.

## Methods

### Participants

Participant number per experiment was constrained to 18 in order to ensure complete counterbalancing of stimulus presentation across participants (explained in details in **Supplementary Note 3**). For Experiment 1, twenty-three healthy participants underwent a pre-selection session (explained in details in **Supplementary Note 4**). This allowed excluding subjects whose overall performance was either at chance or at ceiling with our masked stimuli. Only fifteen participants (10 females and 5 males) matched our selection criterion and were included in the main experiment. Due to external constraints, further recruitment could not be achieved. They were aged 23 ± 3.3. For Experiment 2, 23 participants were recruited, out of which 5 were excluded prior to analysis because their performance was at chance or they did not follow the instructions. The remaining 18 participants (9 females and 9 males) had an average age of 27.5 ± 7.1. In Experiment 3, 26 participants were recruited, out of which 8 were excluded prior to analysis because their overall case discrimination performance was at chance or they did not follow the instructions. Remaining participants (10 females and 8 males) had an average age of 24.5 ± 3.7. Finally, Experiments 4, 5, 6 and 7 included 18 participants each (mean age of the total of 72 participants across studies: 25.4 ± 4.2, 17 men). All participants gave informed consent in writing prior to participation, and the Université Paris Descartes Review Board CERES approved the protocols for the study in accordance with French regulations and the Declaration of Helsinki. Participants received a compensation of 10€ per hour of their time.

### Stimulus-list

Our stimulus list was composed of 432 unique words (targets) from the French language. These words ranged from 3 to 8 letters long (+ one 2-letter word, “os” which translates to “bone”), and did not carry accents of cedillas. They were selected for being strongly associated to another word from the French language less than three-syllables-long, which could then be used as a congruent cue for that word. These associations were determined through 5 different free-association experiments: two former experiments by other groups^70,71^, the online database *dictionnaire des associations verbales du français*^72^ (www.dictaverf.nsu.ru), and two additional online experiments conducted for the purpose of these studies which used the same methodology as the other databases (**Supplementary Note 5**). Cue-target pairs were considered “strongly associated” when, in one free-association experiment, more than 25% of participants first responded the target when presented with the cue. Each word in the target stimulus list was associated with only one possible congruent cue, and only one possible incongruent cue. Incongruent cues were produced once for all experiments by shuffling the list of congruent cues, and checking the absence of semantic relation with the corresponding target word.

### Experimental design

This experiment followed a 2×4 factorial design: congruency (2 levels) and target duration (4 levels), plus an extra level of stimulus absent trials. Each of these 9 conditions included 48 individual trials, split in 15 or 12 blocks. Participants knew that the target was absent on some trials. Word-lists for all participants were established prior to the experiments so as to avoid or balance cross-trial priming and prior exposure effects (**See Supplementary Note 3 & 4** for Participant Selection and Training & Stimulus Presentation Balance).

### Procedure

In Experiment 1, each trial began with the onset of a circle around the central point, when fixation was successfully maintained for 200ms using an EyeTracker (see **Supplementary Note 6** for Materials). Then, following a 1-2s jittered delay, 6 random strings of non-alphanumerical characters (picked randomly without replacement among the following: *%#?&/!|±¶*) with the same font and size as target words would rapidly flash on screen with no inter-stimulus interval. These strings would vary both in length (3 to 8 items, all six possible length being displayed once in each trial) and duration (two masks of each possible duration: 12, 96 or 192ms). We used 6 masks so that all possible target word lengths were presented before presentation of the actual target. This was designed to avoid easily detectible variations in image contrast correlated to target word length. A 7^th^ non-alphanumerical mask would follow suit, be 9-characters long (1 character longer than the target to ensure masking it entirely) and last for 200ms immediately before target onset. Targets would be displayed for 12, 24, 36 or 48ms. There were 11% of “target absent” trials where a mask was presented instead of a target, with the same length and duration probability. The target was then followed by an 8^th^ and last mask that was again 9-characters long and lasted for 200ms. All the sequence unfolded with no blank interval between the stimuli. Fifteen milliseconds after the offset of the last mask, i.e. 215ms after the offset of the visual word target, an auditory cue was presented to the participant. The auditory cue varied in duration depending on word length, and was buffered with silence at the end if necessary for the interval between cue onset and the following event to last for 800ms. It was either semantically related to the target (congruent trials), or unrelated (incongruent trials). All these audio words were pre-recorded with the same female voice and a neutral tone. Participants waited until the offset of the cue (800ms) before they could respond. They first named the target visual word aloud and their response was recorded via a microphone. If they had not seen any word, they were instructed to provide a random word to the best of their ability. Following this first response, participants had to press the space bar or wait for the end of the 10 second recording delay. They were then prompted to report whether the word was written in upper or lower case (and guess if they did not know) by selecting an option on the screen using the keyboard. Finally they had to rate the word’s visibility on a 9-point horizontal scale (a small filled rectangle could be moved within a larger box with ends flanked with the labels “min” and “max”, respectively to the left and the right. See **Supplementary Note 1**). Participants were instructed to use the lowest rating when they had not seen the stimulus at all, and to rate overall visibility using the rest of the scale if they had.

In Experiment 2, the procedure was identical to Experiment 1 except for the following: two synchronous streams of masks were presented just above and below a small fixation circle. Simultaneous masks were different strings of symbols of different lengths that lasted the same duration. On each trial a single target word appeared, in one of the streams, while on the other stream another mask of the same length would be displayed. Note that while there is a known preference for reading in the right visual hemifield^73^ compared to the left, no such strong preference seems to exist between the upper and lower hemifields^74^. In 11% of trials, no target word was presented and the trial only consisted in two streams of masks. While in Experiment 1 the interval between the target offset and the cue onset (ISI) was kept constant at 215ms, in Experiment 2 and 3 it was the interval between target onset and cue onset (SOA) that was kept constant at 248ms, with the introduction of a blank period between last mask offset and cue onset with a duration adjusted according to target duration (between 0 and 36ms). Immediately after the offset of the auditory cue, participants were prompted by a response screen to report where the target had appeared (top or bottom). They then had to provide a visibility rating on this feature using a cursor on a 9-point visual scale similar to Exp. 1 taking into account both detection and position discrimination (**Supplementary Note 1**). Then they were required to report the target word verbally (recorded via a microphone), and then provide a visibility rating taking into account both detection and identification.

Experiment 3 was a follow-up from Experiment 1 in which we changed the order of the tasks (to check for potential task order effects) and included confidence ratings on the two tasks, instead of visibility ratings. The sizes of the stimuli were similar to the ones used in Experiment 2, but they were centered, as in Experiment 1. Participants were first prompted to report whether the target was uppercase or lowercase, and how confident they were about this decision on a 9-point scale. Then they were required to report the target word verbally, and rate their confidence on their word identification response on a 9-point scale. This time, the scales were used to measure only the confidence on the previous decision; there was no mention of visibility. Participants were asked to use the lowest level when they were entirely not-confident about their decision i.e. response at random, the first half of the scale if their confidence was low to medium, and the second half if they were fairly to definitely certain of their response.

In Experiments 4-7, the general structure of the design was kept. 18 participants were included in each according to the criteria listed above, none undertook more than one experiment. The SOA in pre-cueing was 1.6s with the cue appearing before the series of masks, whereas in retro-cueing, the SOA was 248ms. Congruent cues were audio tracks of the same word as the target, neutral cues were the same audio tracks reversed (4&5) or pure tones with the same envelope (6&7). All tracks were tested by three independent reviewers to verify they did not resemble French words. The rating scales instructions were provided as in Experiment 2.

### Data preprocessing

Trial exclusion criteria: For Experiments 1, 2 & 3, trials were excluded when the verbal response was null or inconsistent with instructions, or if expected timings during the trials were not respected. Word identification was performed orally to avoid spelling mistakes had they used the keyboard, and guessing with the help of the auditory cue had they been presented with multiple choices. Audio tracks from each participant were transcribed manually. Trials were rejected when participants were unintelligible, didn’t provide a verbal response, or responded a variation of “I don’t know”, which they were prompted never to say. The average proportion of rejected trials in total was 4 ± 8% (including one outlying participant with 31% rejected trials, who did not comply with the instruction of always providing an answer at the end of a trial. All analyses were also conducted without said outlier, but led to the same conclusions: they were kept for analyses), 3.7± 2.8 % in Experiment 2, and 1.1 ± 0.9 % in Experiment 3, mostly due to participants not responding on some trials. No other exclusions were made on the basis of response times, as time-order of tasks varied in these protocols and we preferred keeping a constant analysis plan rather than creating individual exclusion criteria on this basis. In Experiments 4-7 however, since probes occurred in the same order without verbal responses, we could exclude trials in which participants took over 2.5 median absolute deviation from the median response time to answer under the assumption these trials corresponded to lapses in attention. This led to the exclusion of < 5% of trials in all participants.

### Modeling of word identification and case (or position) discrimination

We analyzed participants’ accuracies at identifying the masked word and discriminating its visual features (e.g., case or position) with a logistic regression^38^. Given that our dataset is grouped according to ‘observational units’ (participants) which could be considered as random sample from a population, we adopted a hierarchical Bayesian approach. For word identification, the probability of identifying the word during a given trial is a mixture of true hits and guessed responses (heard “hive”, tried “bee” without having seen the word). In congruent trials, these guesses correspond to the correct answer and inflates performance, but not in incongruent trials. Under the assumption that participants are about as likely to guess in congruent and incongruent trials when visual information is scarce, we identified guess trials in the incongruent condition based on whether participants had reported the word associated with the cue, and treated them as correct answers when modeling. The logistic regression used was of the form:

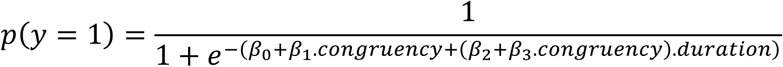

where p(y = 1) corresponds to the probability of correct responses, “duration” corresponds to the duration of the target, and “congruency” is either 0, for incongruent cues, or 1 for congruent cues. This corresponds to a standard logistic regression model estimating the effect of congruency, target duration, and their interaction while including an intercept to capture the estimated guess rate at target duration 0. We also included subject specific random effects for all β parameters as well as word-specific random effects for the intercept and target duration, assuming that words of varying length and lexical frequency could be more or less difficult to identify with little visual information.

All β coefficients in the model were given standard priors with help of the R package Rstanarm^75^. We noted that the proportion of guesses decreases in incongruent trials as the duration of the visual target increases: when more information about the visual target is made available to the participant thanks to longer target durations, it becomes less likely that they will guess using the information of the cue alone. The guess rate estimated from incongruent trials was fitted using a logistic regression estimating the effect of target duration and an intercept on the probability of guess in incongruent trials (guesses in congruent trials are mixed with true hits and cannot be identified), with participant-specific random effects for both coefficients. Using a likelihood-ratio test^76^ we assessed that including target duration significantly improved the model fit vs. an intercept-only model in all experiments (Exp. 1: χ²(1) = 66.488, p <0.001, Exp. 2: χ²(1) = 11.336, p <0.001, Exp. 3: χ²(1) = 49.85, p <0.001). This is shown in **Supplementary Figure 1**.

For visual representations of performance corrected for guess-rate in the figures throughout the article, we subtracted, separately for each target duration, the estimated guess rate in incongruent trials (% of trials where they reported the word associated with the cue) from mean data and estimated performance by the model.

The same model and procedure was used for case or position discrimination performance, except for the correction for guesses which was irrelevant in these cases. For Experiments 4 through 7, the same approach was used but also included type of cue (pre– or retro-) and all possible interactions between the three main effects (duration, congruency, type of cue).

### Model fitting

The model was implemented in Stan^77^ with the help of the Rstanarm package^75^. Markov Chain Monte Carlo (MCMC) sampling was used to draw samples from posterior distribution of model parameters. The model was fitted separately for each experiment, quality of fit was assessed by running 4 sampling MCMC chains and comparing the variance across and within chains with 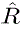. In all models fitted, all 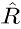 statistics were < 1.01, indicating appropriate convergence (see **Supplementary Table 3** for diagnostic tables of all model fits). We derived predictions for the expected effect of congruency by marginalizing over the distribution of word-specific random effects.

### Detection sensitivity, confidence sensitivity and frequentist analyses

Detection sensitivity and confidence sensitivity were derived from the distributions of visibility or confidence ratings respectively using Signal Detection Theory^41,78^. Areas under the receiver operating characteristic curve (AROC) were computed by comparing distributions of ratings on target-present trials in each considered condition to ratings provided in target-absent trials, using the following relationship:

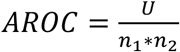

Where U is the result of a Wilcoxon signed-rank test on two independent samples of respective sizes n_1_ and n_2_. These AROC measures were then analyzed using frequentist statistics. Frequentist statistics were also used in the conditional probability analysis described below. The full distributions of visibility and confidence ratings for all conditions can be found in supplementary material.

### Conditional probability analysis: estimating case or position performance in trials with correct word identification

For the conditional performance panels, performance at the case or position task was calculated for trials where the word was correctly identified. In the congruent condition this required discarding “correct-guesses” trials in the congruent condition where they cannot be easily identified. We calculated an individual, per target duration, visibility cut-off below which a correct trial could safely be classified as guess (since the response was correct but the subjective rating was low). If a participant’s point-estimate of the guess rate (based on the guesses identified in the incongruent condition) was n% of congruent trials, we excluded all trials with visibility equal or below that of the n^th^ percentile. This cut-off was most of the time between visibility 0 and visibility 1 (0.27±0.46 for 12ms, 0.47±0.64 for 24ms, 0.74±0.88 for 36ms, 0.87±0.64 for 48ms for Experiment 1).

### Conditional probability analysis: estimating case or position performance in retro-perceived trials

Conditional performance for “retro-perceived” trials, i.e. the additional “word-correct” trials brought about by congruent retro-cues was computed as follows. **Supplementary Figure 3** shows the relationship between different types of events in our experiment in the form of a Venn’s diagram. The totality of trials in the experiment is *C* ∪ *I*, such that *p*(*C* ∪ *I*) = 1, and *p*(*C*) = *p*(*I*) = 0.5, with C and I respectively being the proportion of congruent and incongruent trials. All the quantities that do not involve *R* (i.e. the proportion of retro-perceived words) can be directly calculated from the data.

Note that the figure is not drawn to scale: areas in the diagram do not correspond to the realistic proportions in the data.

We define “retro-perception trials” as follows: retro-perception trials are trials where (i) the word was correctly identified, and (ii) the cue was congruent. This translates as follows in ensemble notation, with W being the proportion of trials with correct word-identification:

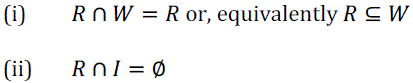

Additionally, retro-perception trials are the trials where congruent cues introduced a difference in performance compared to incongruent cues. This has two implications: (iii) after excluding retro-perceived trials, word-identification performance is similar in incongruent and congruent trials and (iv) after excluding retro-perceived trials, performance at feature identification is similar among word-correct trials *W* in incongruent and congruent trials.

This translates as follows in ensemble notation, with F being the proportion of trials with correct feature-identification:

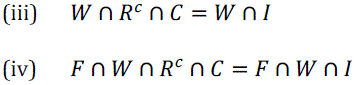

where *R*^c^ indicates the complement of *R* (proportion of trials that are not R).

Within this framework, we calculated the probability of accurately reporting the feature in retro-perception trials. This probability can be expressed as follows:

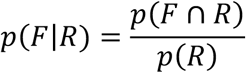

The denominator *p*(*R*) can be obtained as follows:

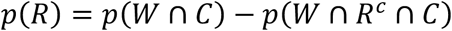

Given (iii) this can be rewritten as follow:

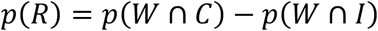

In other words, the probability of retro-perceiving is the difference between the probability of a correct word identification in congruent and incongruent trials.

Similarly, the numerator *p*(*F* ∩ *R*) can be expressed as:

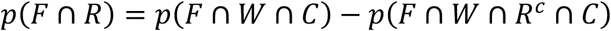

Given (iv), this can be rewritten as follows:

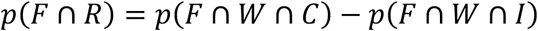

Therefore:

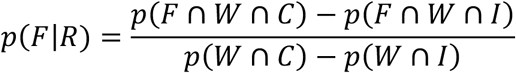

which can be directly calculated from the data.

### Levenshtein distance analysis

To compute string distance in partial awareness controls, we calculated the Levenshtein distance by means of the Wagner-Fischer algorithm^79^. This distance between a string A and a string B is the number of substitutions, insertions or deletions of individual letters needed to turn string A into string B. We used this metric because it is more specific than position-independent letter-matching (is there an “a” in A and B), and less specific than position-dependent letter matching (is there an “a” in position 2 in strings A and B), as it accounts for position in-word (it accounts for fragments of more than one letter, e.g. bigrams or trigrams) but allows for character insertion and deletion. This distance was then divided by the total number of letters in the target word, to obtain reasonably comparable measures.

### Data and script availability

data and scripts are available at https://osf.io/kd2wh/, https://osf.io/t6hxb and https://osf.io/sqcpm as well as from the corresponding author upon reasonable request.

## Acknowledgements

We thank Benjamin Rohaut, Lionel Naccache, Benoît Laslier, Matthias Michel, and Laurent Cohen for their useful comments.

## References

1. Aru, J., Bachmann, T., Singer, W. & Melloni, L. Distilling the neural correlates of consciousness. Neurosci Biobehav Rev 36, 737–46 (2012).

2. Sergent, C. & Naccache, L. Imaging neural signatures of consciousness: ‘What’, ‘When’, ‘Where’ and ‘How’ does it work? Archives italiennes de biologie 150, 91–106 (2012).

3. Hupe, J. M. et al. Cortical feedback improves discrimination between figure and background by V1, V2 and V3 neurons. Nature 394, 784–7 (1998).

4. Lamme, V. A. F. & Roelfsema, P. R. The distinct modes of vision offered by feedforward and recurrent processing. Trends in Neurosciences 23, 571–579 (2000).

5. Bullier, J. Integrated model of visual processing. Brain Res Brain Res Rev 36, 96–107 (2001).

6. Bullier, J., Hupe, J. M., James, A. C. & Girard, P. The role of feedback connections in shaping the responses of visual cortical neurons. Prog Brain Res 134, 193–204 (2001).

7. Super, H., Spekreijse, H. & Lamme, V. A. Two distinct modes of sensory processing observed in monkey primary visual cortex (V1). Nat Neurosci 4, 304–10 (2001).

8. Goldman-Rakic, P. S. Topography of cognition: Parallel distributed networks in primate association cortex. Annual Review of Neuroscience 11, 137–156 (1988).

9. Dehaene, S., Changeux, J.-P., Naccache, L., Sackur, J. & Sergent, C. Conscious, preconscious, and subliminal processing: a testable taxonomy. Trends Cogn. Sci. (Regul. Ed.) 10, 204–211 (2006).

10. Lamme, V. A. Towards a true neural stance on consciousness. Trends in cognitive sciences 10, 494–501 (2006).

11. Tononi, G. Integrated information theory of consciousness: an updated account. Archives italiennes de biologie 150, 56–90 (2012).

12. Boly, M. et al. Are the Neural Correlates of Consciousness in the Front or in the Back of the Cerebral Cortex? Clinical and Neuroimaging Evidence. J Neurosci 37, 9603–9613 (2017).

13. Vandenbroucke, A. R. E., Fahrenfort, J. J., Sligte, I. G. & Lamme, V. A. F. Seeing without Knowing: Neural Signatures of Perceptual Inference in the Absence of Report. Journal of Cognitive Neuroscience 26, 955–969 (2014).

14. de Gardelle, V., Kouider, S. & Sackur, J. An oblique illusion modulated by visibility: Non-monotonic sensory integration in orientation processing. Journal of Vision 10, 6 (2010).

15. de Lafuente, V. & Romo, R. Neural correlate of subjective sensory experience gradually builds up across cortical areas. Proceedings of the National Academy of Sciences 103, 14266–14271 (2006).

16. Lamme, V. A. Why visual attention and awareness are different. Trends Cogn Sci 7, 12–18 (2003).

17. Baars, B. J. Global workspace theory of consciousness: toward a cognitive neuroscience of human experience. in Progress in Brain Research (ed. Laureys, S.) vol. 150 45–53 (Elsevier, 2005).

18. Lau, H. & Rosenthal, D. Empirical support for higher-order theories of conscious awareness. Trends in cognitive sciences 15, 365–373 (2011).

19. Gaillard, R. et al. Converging intracranial markers of conscious access. PLoS Biol 7, e61 (2009).

20. Pitts, M. A., Padwal, J., Fennelly, D., Martínez, A. & Hillyard, S. A. Gamma band activity and the P3 reflect post-perceptual processes, not visual awareness. Neuroimage 101, 337–350 (2014).

21. Fisch, L. et al. Neural “ignition”: enhanced activation linked to perceptual awareness in human ventral stream visual cortex. Neuron 64, 562–574 (2009).

22. Sergent, C., Baillet, S. & Dehaene, S. Timing of the brain events underlying access to consciousness during the attentional blink. Nature neuroscience 8, 1391 (2005).

23. Hesselmann, G., Kell, C. A., Eger, E. & Kleinschmidt, A. Spontaneous local variations in ongoing neural activity bias perceptual decisions. Proc Natl Acad Sci U S A 105, 10984–9 (2008).

24. Romei, V. et al. Spontaneous Fluctuations in Posterior α-Band EEG Activity Reflect Variability in Excitability of Human Visual Areas. Cerebral Cortex 18, 2010–2018 (2008).

25. Sergent, C. The offline stream of conscious representations. Philosophical Transactions of the Royal Society B: Biological Sciences 373, 20170349 (2018).

26. Rimsky-Robert, D., Störmer, V., Sackur, J. & Sergent, C. Retrospective auditory cues can improve detection of near-threshold visual targets. Scientific reports 9, 1–11 (2019).

27. Garnier-Allain, A., Pressnitzer, D. & Sergent, C. Retrospective cueing mediates flexible conscious access to past spoken words. Journal of Experimental Psychology: Human Perception and Performance (accepted pending minor revisions) (2023).

28. Thibault, L., Van den Berg, R., Cavanagh, P. & Sergent, C. Retrospective attention gates discrete conscious access to past sensory stimuli. PloS one 11, e0148504 (2016).

29. Xia, Y., Morimoto, Y. & Noguchi, Y. Retrospective triggering of conscious perception by an interstimulus interaction. Journal of vision 16, 3–3 (2016).

30. Enns, J. T. & Di Lollo, V. What’s new in visual masking? Trends in Cognitive Sciences 4, 345–352 (2000).

31. Dehaene, S. et al. Imaging unconscious semantic priming. Nature 395, 597 (1998).

32. Kiefer, M. & Spitzer, M. Time course of conscious and unconscious semantic brain activation. NeuroReport 11, 2401 (2000).

33. Kouider, S. & Dehaene, S. Levels of processing during non-conscious perception: a critical review of visual masking. Philos Trans R Soc Lond B Biol Sci 857–875 (2007).

34. Collins, A. M. & Loftus, E. F. A spreading-activation theory of semantic processing. Psychological review 82, 407 (1975).

35. Sandberg, K., Timmermans, B., Overgaard, M. & Cleeremans, A. Measuring consciousness: is one measure better than the other? Consciousness and Cognition 19, 1069–78 (2010).

36. Sergent, C. & Dehaene, S. Is consciousness a gradual phenomenon? Evidence for an all-or-none bifurcation during the Attentional Blink. Psychol Sci 15, 720–728 (2004).

37. Garnier-Allain, A., Pressnitzer, D. & Sergent, C. Retrospective cueing mediates flexible conscious access to past spoken words. J Exp Psychol Hum Percept Perform 49, 949–967 (2023).

38. Cox, D. R. The analysis of multivariate binary data. Applied statistics 113–120 (1972).

39. Bakeman, R. Recommended effect size statistics for repeated measures designs. Behavior Research Methods 37, 379–384 (2005).

40. Cohen, J. The effect size. Statistical power analysis for the behavioral sciences 77–83 (1988).

41. Hanley, J. A. & McNeil, B. J. The meaning and use of the area under a receiver operating characteristic (ROC) curve. Radiology 143, 29–36 (1982).

42. Kouider, S., De Gardelle, V., Sackur, J. & Dupoux, E. How rich is consciousness? The partial awareness hypothesis. Trends in cognitive sciences 14, 301–307 (2010).

43. Kouider, S., de Gardelle, V. & Dupoux, E. Partial awareness and the illusion of phenomenal consciousness. Behavioral and Brain Sciences 30, 510–511 (2007).

44. Kouider, S. & Dupoux, E. Partial awareness creates the “illusion” of subliminal semantic priming. Psychological science 15, 75–81 (2004).

45. Dark, V. J. Semantic priming, prime reportability, and retroactive priming are interdependent. Memory & Cognition 16, 299–308 (1988).

46. VanVoorhis, B. A. & Dark, V. J. Semantic matching, response mode, and response mapping as contributors to retroactive and proactive priming. *Journal of Experimental Psychology: Learning*, Memory, and Cognition 21, 913 (1995).

47. Bernstein, I. H., Bissonnette, V., Vyas, A. & Barclay, P. Semantic priming: Subliminal perception or context? Perception & Psychophysics 45, 153–161 (1989).

48. Neely, J. H. Semantic priming effects in visual word recognition: A selective review of current findings and theories. Basic processes in reading: Visual word recognition 11, 264–336 (1991).

49. Briand, K., Heyer, K. den & Dannenbring, G. L. Retroactive semantic priming in a lexical decision task. The Quarterly Journal of Experimental Psychology 40, 341–359 (1988).

50. Thomas, M. A., Neely, J. H. & O’Connor, P. When word identification gets tough, retrospective semantic processing comes to the rescue. Journal of Memory and Language 66, 623–643 (2012).

51. Seth, A. K., Dienes, Z., Cleeremans, A., Overgaard, M. & Pessoa, L. Measuring consciousness: relating behavioural and neurophysiological approaches. Trends in cognitive sciences 12, 314–321 (2008).

52. Derrien, D., Garric, C., Sergent, C. & Chokron, S. The nature of blindsight: implications for current theories of consciousness. Neuroscience of Consciousness 2022, niab043 (2022).

53. Weiskrantz, L. Consciousness Lost and Found: A Neuropsychological Exploration. (Oxford University Press, New York, 1997).

54. Stiles, N. R., Li, M., Levitan, C. A., Kamitani, Y. & Shimojo, S. What you saw is what you will hear: Two new illusions with audiovisual postdictive effects. PloS one 13, e0204217 (2018).

55. Maynes, R. et al. Metacognitive awareness in the sound-induced flash illusion. Phil. Trans. R. Soc. B 378, 20220347 (2023).

56. Lamme, V. A. F., Zipser, K. & Spekreijse, H. Masking Interrupts Figure-Ground Signals in V1. Journal of Cognitive Neuroscience 14, 1044–1053 (2002).

57. Dehaene, S. et al. Cerebral mechanisms of word masking and unconscious repetition priming. Nature neuroscience 4, 752 (2001).

58. Kouider, S., de Gardelle, V., Dehaene, S., Dupoux, E. & Pallier, C. Cerebral bases of subliminal speech priming. Neuroimage 49, 922–929 (2010).

59. Sprague, T. C., Ester, E. F. & Serences, J. T. Restoring Latent Visual Working Memory Representations in Human Cortex. Neuron 91, 694–707 (2016).

60. Stokes, M. G. ‘Activity-silent’ working memory in prefrontal cortex: a dynamic coding framework. Trends Cogn Sci 19, 394–405 (2015).

61. Harrison, S. A. & Tong, F. Decoding reveals the contents of visual working memory in early visual areas. Nature 458, 632–5 (2009).

62. Serences, J. T., Ester, E. F., Vogel, E. K. & Awh, E. Stimulus-specific delay activity in human primary visual cortex. Psychol Sci 20, 207–14 (2009).

63. Lamme, V. A., Super, H., Landman, R., Roelfsema, P. R. & Spekreijse, H. The role of primary visual cortex (V1) in visual awareness. Vision research 40, 1507–1521 (2000).

64. Sergent, C. The offline stream of conscious representations. Philosophical Transactions of the Royal Society B: Biological Sciences 373, 20170349 (2018).

65. Sergent, C. et al. Cueing attention after the stimulus is gone can retrospectively trigger conscious perception. Current Biology 23, 150–155 (2013).

66. Faw, B. Conflicting Intuitions May Be Based On Differing Abilities: Evidence from Mental Imaging Research. Journal of Consciousness Studies 16, 45–68 (2009).

67. Hochstein, S. & Ahissar, M. View from the top: Hierarchies and reverse hierarchies in the visual system. Neuron 36, 791–804 (2002).

68. Summerfield, C. & de Lange, F. P. Expectation in perceptual decision making: neural and computational mechanisms. Nature Reviews Neuroscience 15, 745–756 (2014).

69. Lupyan, G. & Ward, E. J. Language can boost otherwise unseen objects into visual awareness. Proceedings of the National Academy of Sciences 110, 14196–14201 (2013).

70. Ferrand, L. & Alario, F.-X. Normes d’associations verbales pour 366 noms d’objets concrets. L’Année psychologique 98, 659–709 (1998).

71. Ferrand, L. Normes d’associations verbales pour 260 mots «abstraits». L’Année psychologique 101, 683–721 (2001).

72. Debrenne, M. Le dictionnaire des associations verbales du français et ses applications. Variétés, variations and formes du français. Palaiseau: Éditions de l’Ecole polytechnique 355–66 (2011).

73. Ellis, A. W. Length, formats, neighbours, hemispheres, and the processing of words presented laterally or at fixation. Brain and language 88, 355–366 (2004).

74. Hagenbeek, R. E. & Van Strien, J. W. Left–right and upper–lower visual field asymmetries for face matching, letter naming, and lexical decision. Brain and Cognition 49, 34–44 (2002).

75. Goodrich, B., Gabry, J., Ali, I. & Brilleman, S. rstanarm: Bayesian applied regression modeling via Stan. R package version 2, (2020).

76. Pinheiro, J. C. & Bates, D. M. Linear mixed-effects models: basic concepts and examples. Mixed-effects models in S and S-Plus 3–56 (2000).

77. Carpenter, B. et al. Stan: A probabilistic programming language. Journal of statistical software 76, 1–32 (2017).

78. McMillan, N. A. & Creelman, C. D. Detection theory. LEA Pub (2005).

79. Wagner, R. A. & Fischer, M. J. The string-to-string correction problem. Journal of the ACM (JACM) 21, 168–173 (1974).

