## Supplementary Information for "Consciously detecting and recognizing a past visual word after its sensory trace is gone"

### Supplementary Notes

**Supplementary Note 1: Instructions.**

Here are the general instructions provided to the participants in the various experiments (instructions relative to Experiment 2 are in between parentheses):

*On each experimental trial, you will see a circle and a dot in its center. You will have to fixate this circle. The eyetracker follows your gaze, and waits until you are fixating to start. When the trial starts, you will see character strings appear in the center of the screen (expt2: above and below the center). They will make no sense at all. Among them, most of the time, there will be a meaningful word written in uppercase or lowercase (expt2: in uppercase), but in about 10% of trials, there will be no word at all. (Please make sure to fixate the central dot, otherwise you will not be able to read properly.) After this visual sequence, you will hear another, different word in the headset you are wearing. This word will be related to the written word half the time e.g. witch, broom. This word can help you, or have no effect at all, try and pay attention to it.*

The task instructions for Experiment 1 were:

*You will answer 3 questions:*

1) *What was the written word among the nonsensical strings? If you do not know, say whatever comes to your mind. You should always provide an answer, and try and say just one word. If you're having difficulties coming up with different words every time, you can say the same one when you really don't know, but we encourage you to be spontaneous, it's more fun. "I don't know" is not a valid answer.*
2) *Was this word in uppercase or lowercase?*
3) *How well did you see this word? You have a scale ranging from 0 to 8, choose the appropriate number. 0 is special: it means you have not seen the word at all. 1 corresponds to "maybe a glimpse of something", and 8 means you have perfectly seen the word.*

The task instructions for Experiment 2 were:

*You will answer 4 questions:*

1) *Was the word presented above or below the fixation point?*
2) *How well did you see the word* ***and its position****? You have a colored scale ranging from red to green, with 9 points.*
   *0: I have not seen the stimulus, not even a glimpse of a letter.*

   *1-4: I have seen something, I may have seen a few letters, but I'm not really sure* ***where*** *(focus on position) this was.*

   *5-8: I have seen the word* ***and its position****. Increase with stronger sensation of having seen the word and its position.*

3) *What was the word written among the nonsensical strings? If you don't know, say whatever comes to mind. Always provide an answer, try and say just one word. If you're having difficulty coming up with different words every time, you can say the same one when you really don't know, but we encourage you to be spontaneous, it's more fun. "I don't know" is not a valid answer.*
4) *How well did you see the word* ***and its identity****? You have a colored scale ranging from red to green, with 9 points.*
   *0: I have not seen the stimulus, not even a glimpse of a letter.*

   *1-4: I have seen something, I may have seen a few letters, but I'm not really sure* ***what*** *(focus on identity/reading) this was.*

   *5-8: I have seen the word* ***and its identity****. Increase with stronger sensation of having seen the word and its identity*

The task instructions for Experiment 3 were:

*You will answer 4 questions:*

1) *Was this word in uppercase or lowercase?*

2) *How confident are you in this uppercase/lowercase response? You have a colored scale ranging from red to green, with 9 points. The lowest rating (completely red) corresponds to no confidence at all, and the highest means you are fully confident.*

*0: not confident at all (reported at chance).*

*1-4: I am not very confident that I correctly reported the word's case.*

*5-8: I am more and more confident that I correctly reported the word's case.*

3) *What was the word written among the nonsensical strings? If you don't know, say whatever comes to mind. Always provide an answer, try and say just one word. If you're having difficulty coming up with different words every time, you can say the same one when you really don't know, but we encourage you to be spontaneous, it's more fun. "I don't know" is not a valid answer.*

4) *How confident are you with your word identity response? You have a colored scale ranging from red to green, with 9 points. The lowest rating (completely red) corresponds to no confidence at all, and the highest means you are fully confident.*

*0: not confident at all (reported at chance).*

*1-4: I am not very confident that I correctly reported the word's identity.*

*5-8: I am more and more confident that I correctly reported the word's identity.*

*The two confidence measures are distinct: you may be confident about the word's case because you caught a single letter, but not know what the word was. The reverse is also possible. Freely reply to each rating separately.*

In Experiment 4 & 5, participants received near-identical instructions, however they were not warned that the neutral cues were reversed audio files for the congruent cues. Instructions for use of the visibility scale were as in Experiment 1.

**Supplementary Note 2: Results of post-hoc comparisons in all Experiments**

Experiment 1, word identification (2b): 12ms, 95%BCI-diff [-0.03 0.09]; 24ms, 95%BCI-diff [0.002, 0.16]; 36ms, 95%BCI-diff [0.02, 0.21]; 48ms, 95%BCI-diff [0.02, 0.25].

Experiment 1, detection sensitivity (2d): 12ms: CI = [-0.03, 0.04] $d$ = 0.08, 24ms: CI = [-0.009, 0.08] $d$ =0.51, 36ms: CI = [0.03, 0.08] $d$ = 1.36, 48ms: CI = [0.01, 0.05] $d$ =0.84.

Experiment 2, word identification (3b): 12ms (95%BCI-diff [-0.05 0.04]), 24ms (95%BCI-diff [-0.02 0.06]), 36ms (95%BCI-diff [-0.01 0.12]), 48ms (95%BCI-diff [-0.005 0.18]).

Experiment 2, detection sensitivity word (3d): 12ms (95%CI [-0.07 0.01], $d$ = 0.40), 24ms (95%CI [-0.02 0.04], $d$ = 0.15), 36ms (95%CI [0.02 0.08], $d$ = 0.90), 48ms (95%CI [0.03 0.09], $d$ = 1.03).

Experiment 2, detection sensitivity position (3e): 12ms (95%CI [-0.03 0.03], $d$ = 0.09), 24ms (95%CI [-0.03 0.04], $d$ = 0.01), 36ms (95%CI [0.01 0.07], $d$ = 0.65), 48ms (95%CI [0.01 0.05], $d$ = 0.79).

Experiment 3, word identification (4b): 12ms (95%BCI-diff [-0.05 0.03]), 24ms (95%BCI-diff [-0.003 0.09]), 36ms (95%BCI-diff [0.06 0.18]), 48ms (95%BCI-diff [0.06 0.25]).

Experiment 3, confidence sensitivity word (4d): 12ms (95%CI [-0.03 0.01], $d$ = 0.39), 24ms (95%CI [-0.003 0.6], $d$ = 0.53), 36ms (95%CI [0.01 0.1], $d$ = 0.69), 48ms (95%CI [0.03 0.11], $d$ = 0.87).

Experiment 3, confidence sensitivity case (4e): 12ms (95%CI [-0.01 0.03], $d$ = 0.13), 24ms (95%CI [-0.04 0.02], $d$ = 0.26), 36ms (95%CI [-0.03 0.05], $d$ = 0.16), 48ms (95%CI [0.02 0.08], $d$ = 1.01).

Experiments 4-7, detection sensitivity (5a): 95%CI 12ms [-0.06 0.01], 24ms [-0.0003 0.08], 36ms [0.001 0.07] $d$ = -0.39, 48ms [0.01 0.07] $d$ = -0.51

Experiments 4-7, case performance, pre-cued trials (5b): 12ms (95%BCI-diff [-0.02 0.06]), 24ms (95%BCI-diff [0.01 0.06]), 36ms (95%BCI-diff [0.01 0.06]), and 48ms (95%BCI-diff [0.004 0.04])

Experiments 4-7, case performance, retro-cued trials (5b): 12ms (95%BCI-diff [-0.02 0.05]), 24ms (95%BCI-diff [-0.02 0.03]), 36ms (95%BCI-diff [-0.02 0.03]), and 48ms (95%BCI-diff [-0.03 0.02])

**Supplementary note 3: Stimulus presentation counter-balancing across participants.** For each participant, some of the words in the general list were removed, corresponding to the target absent trials. The words that were removed were counterbalanced across participants so that each target word in the general list would be absent the same number of times across all participants. Second, since each target word was presented only once per participant (to avoid any prior exposure effect), we made sure that, across all participants, each target would appear the same number of times in each of the different conditions. Third, potential priming from one trial to the next by auditory cues was controlled for by having pairs of participants receive the same auditory cues in the same order, but inverting the congruency relationship between the auditory cues and the preceding visual words (e.g. if for the first participant of a pair, a cue followed the associated congruent target word, for the other participant the same cue would follow the incongruent one). Cues could appear from zero to twice (in a congruent and an incongruent trial in this case) for each participant. Incongruent pairs were designed by associating a cue to another target that shared no obvious association to it, and never changed throughout the experiments. In detail, pseudo-randomization was conducted with a focus on balancing cross-trial word priming: the alphabetical stimulus list was shuffled, then divided into 9 blocks of 48 stimuli. For the first pair of participants, the first block was set as target absent trials. For participant 1, pseudorandom (balanced over the entire set) congruency, target duration and feature type was attributed to each non-absent target. Target absent-trials were attributed a pseudorandom target duration. Stimulus-order was then shuffled again to mix target absent trials with target present trials. Participant 2 was presented with the same twice-shuffled stimulus-list, with congruency reversed. The procedure was repeated for 9 participant pairs, so that each of the 9 blocks of stimuli could be used as target absent trials.

**Supplementary Note 4: Participants' pre-selection procedure in Experiment 1.** Given the difficulty of the task, we chose in Experiment 1 to run a pre-selection session that was designed to exclude participants unable to perform the task from the get-go, as well as provide others with some training. We took this opportunity to test a variety of target durations in order to choose the best suited for the testing session. This session lasted 1 hour and included 140 trials preceded by a short, 18 trial training block with progressively decreasing target duration, from 120ms to 12ms. During the preselection session, participants were presented with visually masked words but no auditory cue. The masking procedure was the same as in the main experiment. The target words were different from the main experiment, to avoid prior exposure effects for those participants who were selected, but matched the same criteria. Tested target durations during the preselection session always included 24ms and 36ms, and the 3$^{rd}$ and 4$^{th}$ duration tested ranged from 48ms to 94ms across participants. Participants were asked to identify the word aloud in a microphone, and report its case and visibility using the keyboard. The criterion for inclusion in our experiment was that performance at the case discrimination task should, for at least one target duration, be within our predefined bounds [65% - 90%]. In Experiments 2 & 3, we used a different selection procedure. The experiment was performed over a single session, which started with a 72 trial-long training session with progressively decreasing target duration without any auditory cueing. If participants reported being unable to do the task, and performance suggested as much, we ran a second 72 trial-long training session, at the end of which we evaluated whether a given participant could perform the task.

**Supplementary Note 5: Free association experiments and word selection**. We conducted two online experiments, in which participants were given a single word and prompted to type in the first word that came to mind in response. They were instructed that they had 15 minutes to respond to roughly 100 words. The first experiment tested 203 words and had between 40 and 67 responders to each unit. Data was pre-processed with a semi-automatic spellcheck using the Office suite for Windows. The software detected anomalies, and the experimenter chose the appropriate correction depending on whether the anomaly was an obvious spelling error or not (unintelligible responses were kept as is). Out of these, 137 item pairs fared above the threshold for use on our experiments, which was that >= 25% of responders gave the same answer to one item. The second experiment tested 95 words and had 70 to 79 responders to each of them. Out of these, 73 item pairs were kept according to our criteria. One obvious internet troll was excluded from these experiments, as he responded a single profanity to all proposed items. We otherwise followed the methods of Debrenne et al.[1] Word selection from the online database[1] was performed with our participant demographic in mind: we filtered French native speakers, 18 to 25 years old, and selected words matching our criteria. Extra words were then selected from existing articles on free association[2,3] with the same criteria to complete our list. Words were of all grammatical classes, but kept in dictionary form if possible.

**Supplementary Note 6: Materials**. Materials slightly differed between Experiment 1 and the other two. Differences are indicated in parentheses throughout. Stimuli were generated and responses recorded using the Psychophysics Toolbox for Matlab49. Stimuli were presented on a CRT monitor Sony Trinitron GDM-F520 (Sony MULTISCAN E400 in Exp.2&3). Refresh rate was 85 Hz and screen resolution was 1280 by 1024 pixels (1152 by 864 in Exp.2&3). Participants were seated 70cm away from the monitor (57cm in Exp.2&3), in a dimly lit room. Eye fixation was monitored and recorded using an Eye Tribe tracking device (an EyeLink 1000+ for Exp.2&3). Stimuli were presented on a gray background (52.5 cd/m² in Exp.1; 44.5 cd/m² in Exp.2&3), and participants were told to fixate a black dot inside a small black circle of 1.6° of visual angle in diameter (0.4° in Exp.2&3) at the onset of each trial. In all three experiments, the start of a trial was conditioned on appropriate fixation for at least 200ms (less than 2° from center for Exp.1; criterion was 1.2° for Exp.2&3). Targets were words written in a mono-spaced font (Courier New), either in uppercase or lowercase. They were approximately 1° high and 2° to 7.4° large depending on word-length in Exp.1 (approximately 0.6° high by 1.6° to 5° large in Exp.2&3). Note that the entire display was smaller in Experiment 2 to ensure that all stimuli would still be presented within reading range even though they were displaced from the center of the screen by 0.7°. These changes in size were kept for Experiment 3, as bigger words may be more difficult to read. A HyperX Cloud headset with microphone (Kingston Technology Company Inc.) was used to play the auditory words (retro-cues) and record participants' verbal responses. Audio-visual delay was measured using a photodiode applied to the computer screen and the computer's audio output, together with an oscilloscope for visualization. The delay was within a range of 10ms with no identifiable jitter. This delay was taken into account in the stimulation scripts.

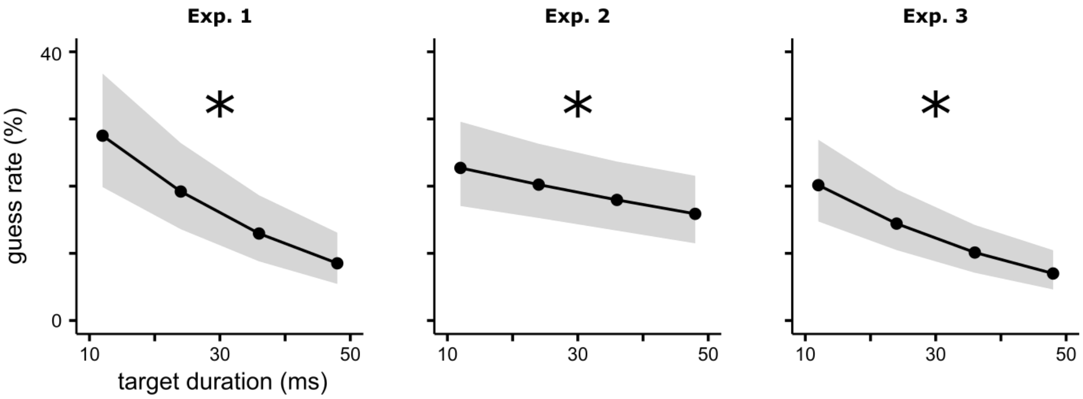


**Supplementary Figure 1: guess rate as a function of target duration.** Guessed trials were defined as trials in the incongruent condition in which participants reported the word associated to the cue (which is as a wrong answer in incongruent trials). A generalized linear mixed-effect model was used to model the guess rate using a logistic regression testing for the fixed-effect of target duration vs. an intercept-only model. The black line represents the posterior estimate of the guess rate, the shaded area is the 95%BCI of the estimated values. Participant number was used as a random-effect on the intercept. Stars denote a significant main effect of target duration.

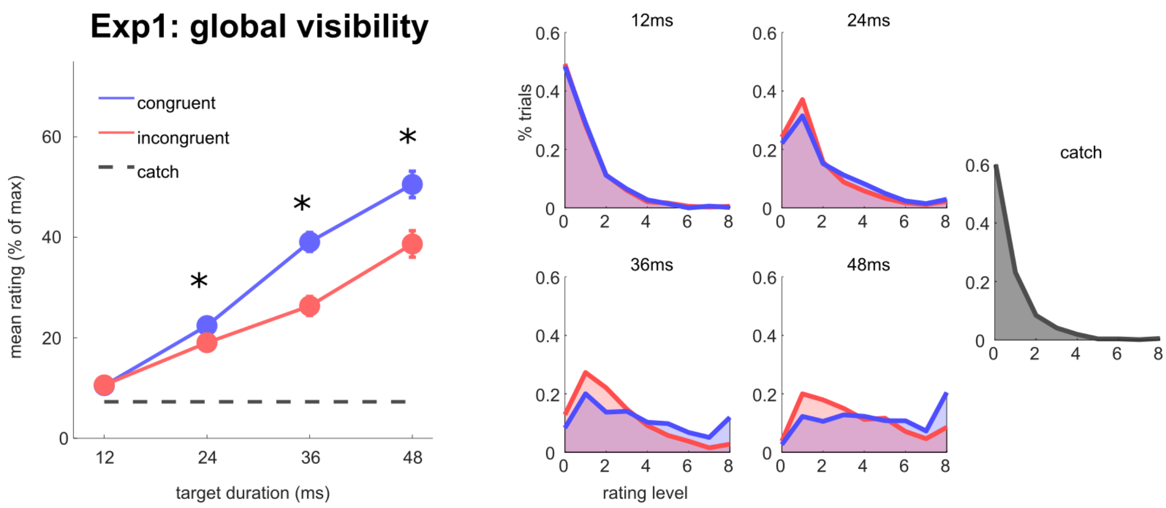


**Supplementary Figure 2: Mean visibility ratings and distributions (Experiment 1).** Left: each dot represents the average across participants of the mean visibility rating obtained on the scale for catch (grey), congruent trials (blue) or incongruent trials (red) as a function of target duration, as well as mean visibility obtained for catch trials (dashed grey). The error bars represent the standard error of the mean difference between congruent and incongruent trials. A repeated measures two-way ANOVA on mean subjective visibility ratings revealed a significant increase in visibility with target duration ($F(1.23,17.29)=81.34$, $p<0.001$) and congruence ($F(1.00,14.00)=38.43$, $p<0.001$), and a significant interaction between these two factors ($F(3.00,42.00)=15.812$, $p<0.001$). Distributions of visibility ratings are shown on the right for each target duration and for catch trials. Notice that even for catch trials, participants sometimes used visibilities higher than 0 to express that they might have felt a glimpse of the stimulus. Detection sensitivity is obtained by comparing each of the distributions to the distribution obtained on catch trials (see Methods).

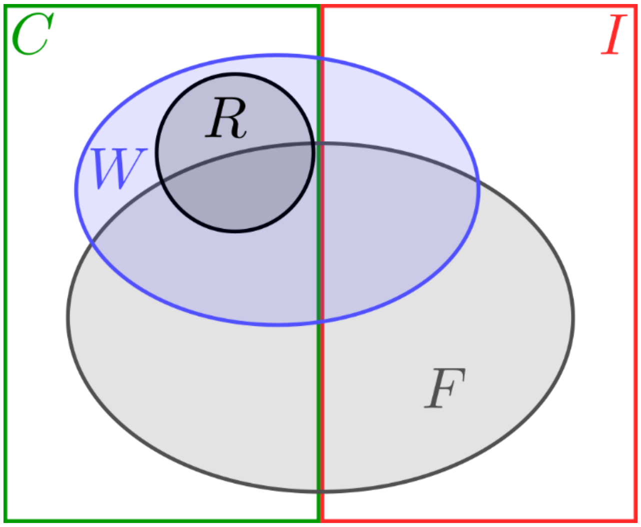


**Supplementary Figure 3: Graphical representation of the probabilities as a Venn diagram.** C, congruent trials; I, incongruent trials; W, trials with correct word-identification; F, trials with correct feature-response (case or position); R, retro-perceived trials, i.e. congruent trials with a correct word-identification response that was obtained via retro-perception.

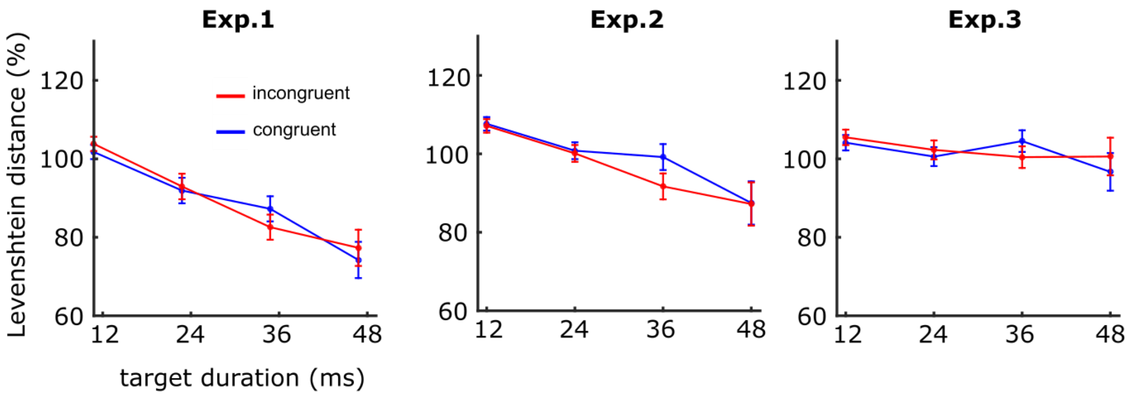


**Supplementary Figure 4: orthographic distance (Levenshtein distance) between erroneous identification responses and the actual target word.** The Levenshtein distance between target and response in word-incorrect trials is shown as a function of target duration and cue congruency. In Experiment 1, the repeated measures ANOVA (duration x congruency) showed an effect of duration ($F(2.45,34.23)=40.157$, $p< 0.01$) but no effect of congruency ($F(1.00,14.00)=0.051$, $p=0.82$) nor an interaction ($F(2.26,31.62)=1.055$, $p=0.37$). In Experiment 2, the repeated measures ANOVA (duration x congruency) showed an effect of duration ($F(2.31,39.32)=3.584$, $p=0.031$) but not effect of congruency ($F(1.00,17.00)=0.209$, $p=0.653$) nor an interaction ($F(2.23,37.93)=1.150$, $p=0.332$). In Experiment 3 the repeated measures ANOVA (duration x congruency) showed an effect of duration ($F(2.17,36.93)=22.790$, $p<0.001$) but not effect of congruency ($F(1.00,17.00)=1.891$, $p=0.187$) nor an interaction ($F(1.68,28.57)=0.960$, $p=0.381$). Note that for some estimates, the number of trials per participant was quite low, especially for longer target durations as only incorrect trials for word identification are used. However, the average number of trials in the 48ms condition (out of a total of 96 trials) was still reasonable in all three experiments: Experiment 1 [43.5], Experiment 2 [63.2], Experiment 3 [42.1]). Error bars represent the standard error of the mean.

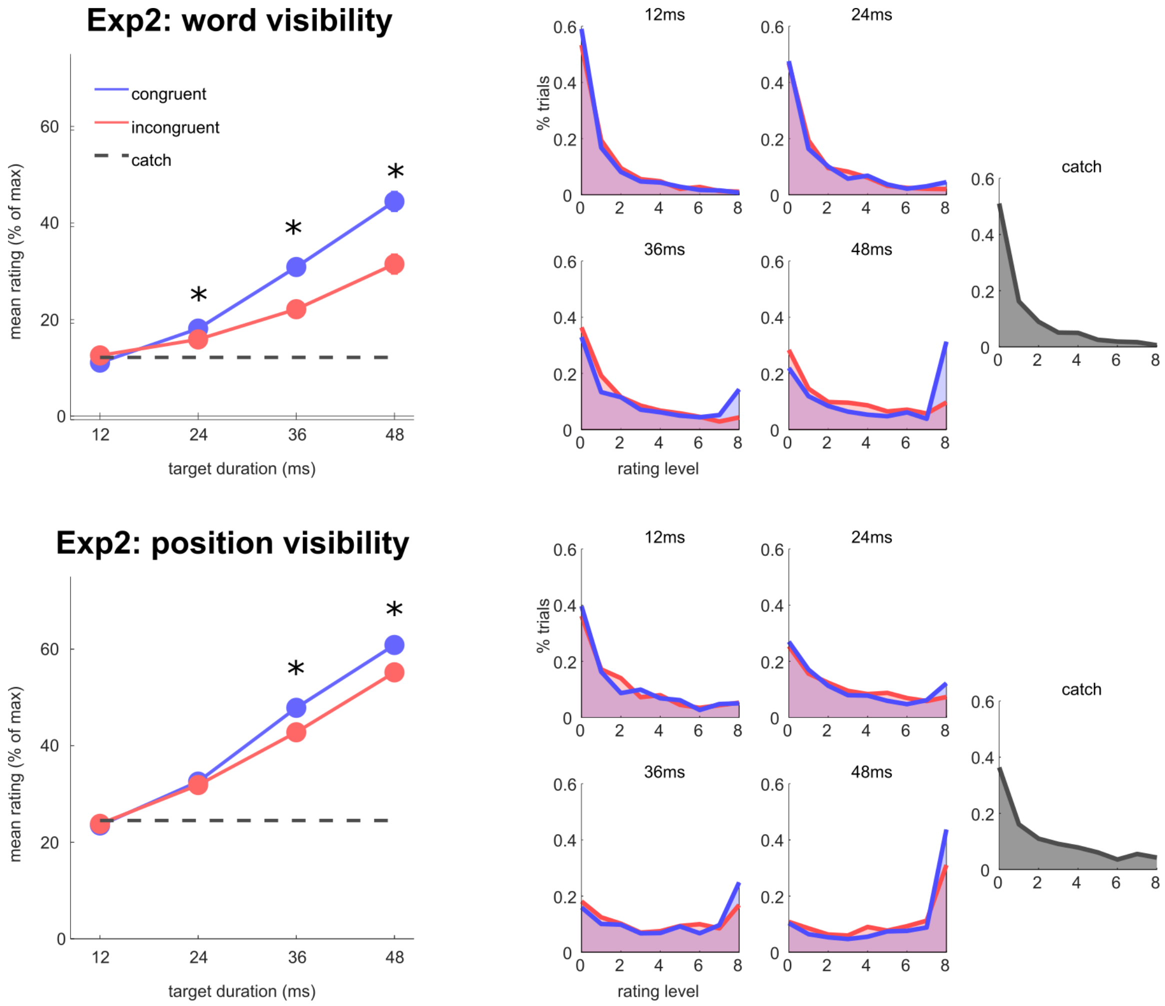


**Supplementary Figure 5: Mean visibility ratings and distributions in Experiment 2, separately for word and position.** In experiment 2, participants rated separately the visibility of the word's identity (upper panel) and position (lower panel). Each dot represents the average across participants of the mean visibility rating obtained for catch (grey), congruent trials (blue) or incongruent trials (red) as a function of target duration. The error bars represent the standard error of the mean difference between congruent and incongruent trials. A repeated measures two-way ANOVA on mean visibility ratings for word-meaning revealed a significant increase with target duration ($F(1.19,20.28)=41.70$, $p<0.001$) and congruence ($F(1.00,17.00)=9.57$, $p<0.001$), with an interaction between these two factors ($F(3.00,51.00)=24.05$, $p<0.001$). A repeated measures two-way ANOVA on mean visibility ratings for position revealed a similar pattern: a significant increase with target duration ($F(1.35,22.94)=64.103$, $p<0.001$) and congruence ($F(1.00,17.00)=30.305$, $p<0.001$), with a significant interaction between these two factors ($F(2.46,41.82)=6.721$, $p=0.002$). Distributions of visibility ratings are shown on the right.

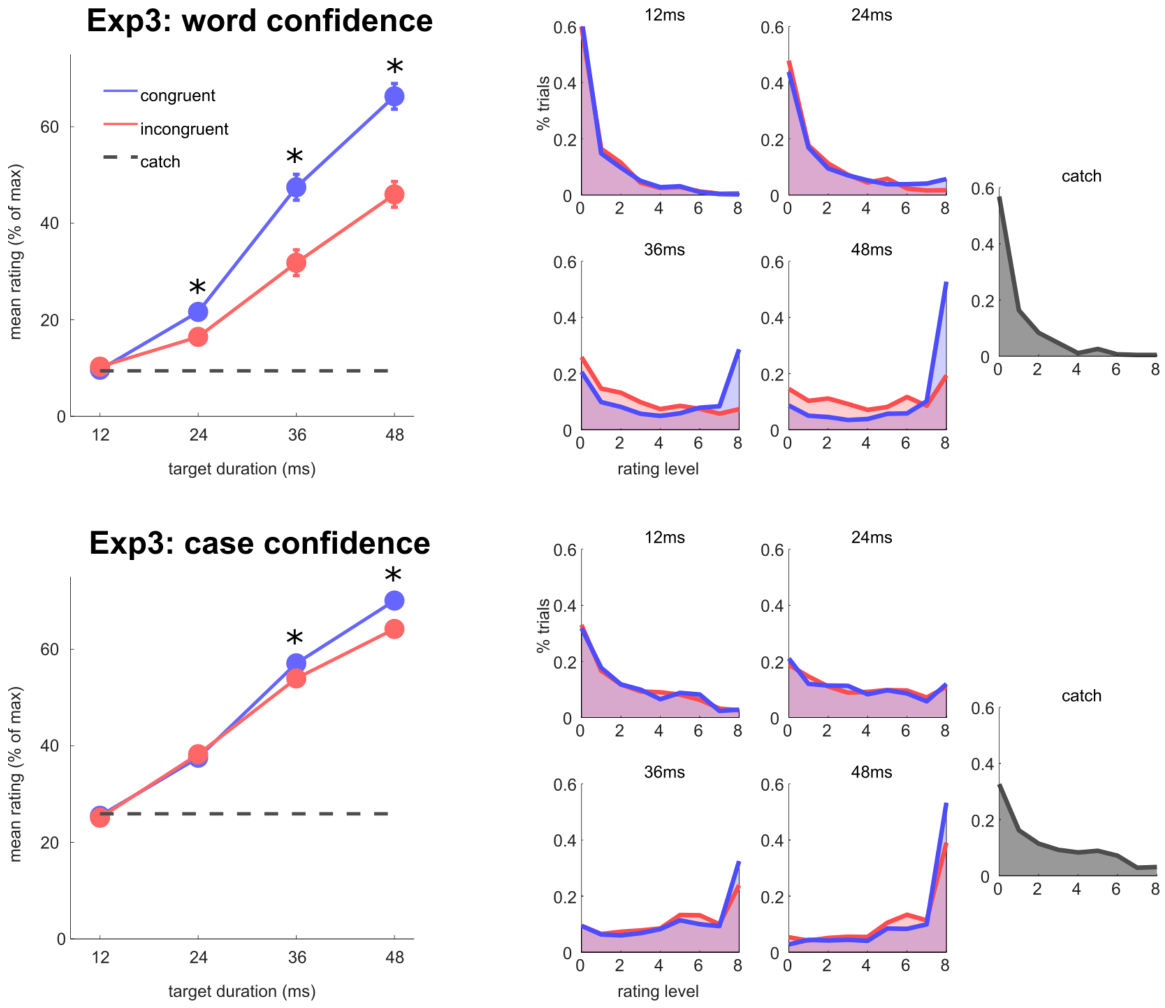


**Supplementary Figure 6: Mean confidence ratings (Experiment 3) and confidence distributions, separately for word identification and case discrimination.** In Experiment 3, participants were requested to rate their confidence on their word identification response (upper panel), and their case discrimination response (lower panel). Each dot represents the average across participants of the mean confidence rating obtained on the scale for catch (grey), congruent trials (blue) or incongruent trials (red) as a function of target duration. The error bars represent the standard error of the mean difference between congruent and incongruent trials. A repeated measures two-way ANOVA on confidence ratings for word identification revealed a significant increase in confidence with target duration ($F(1.24,21.16)=132.037$, $p<0.001$) and congruence ($F(1.00,17.00)=65.538$, $p<0.001$), with an interaction between these two factors ($F(2.12,36.12)=28.223$, $p<0.001$). A repeated measures two-way ANOVA on confidence ratings for case discrimination revealed a significant increase in confidence with target duration ($F(1.31,22.31)=69.95$, $p<0.001$) and congruence ($F(1.00,17.00)=9.07$, $p<0.01$), with an interaction between these two factors ($F(1.31,22.31)=7.05$, $p<0.01$). Distributions of visibility ratings are shown on the right.

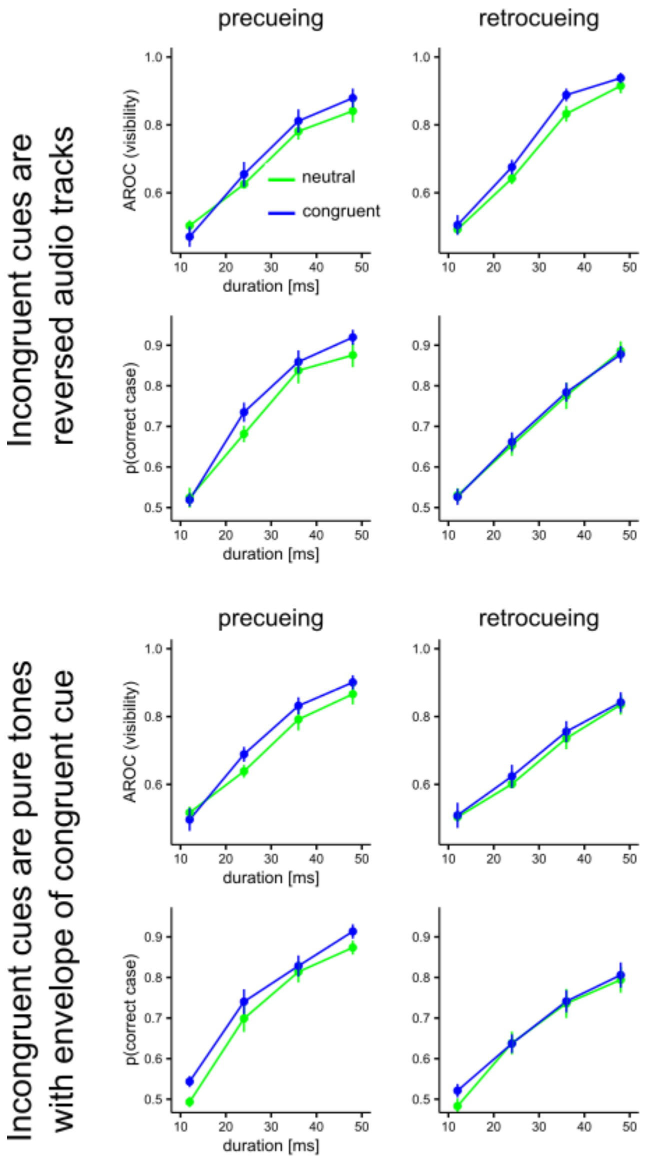


**Supplementary Figure 7: Results for Experiments 4 (precue, reversed track), 5 (retrocue, reversed track), 6 (precue, pure tones), and 7 (retrocue, pure tones) shown as in Figure 5 (where data is pooled 4&6/5&7).** Legend is as in Figure 5.

***Supplementary Table 1: Examples of stimuli. Available in French upon request.*** *Here is a short table showing the first 20 stimulus-pairs from our bank by alphabetical (target) order, with their English translation. The full document is available below (****Supplementary Table 2****). The two first columns correspond to possible congruent and incongruent cues, for the target shown in the third. The last column displays the percentage of association estimated between the target and the corresponding congruent cue as estimates through the procedure described in* ***Supplementary Note 6****. The incongruent pairs are constituted of cue switches (see example highlighted in green).*

**Supplementary Table 2: Full stimulus list. Available in French upon request.**

**Supplementary Table 3: Model fit diagnostics (main effects).**

| | Target duration | congruency | | Interaction dur x congr | | | |
|---|---|---|---|---|---|---|---|
| **Exp 1** | | | | | | | |
| Rhat identification | 1.003 | 1.002 | | 1.001 | | | |
| n_eff identification | 1150 | 3191 | | 2834 | | | |
| Rhat discrimination | 1.003 | 1.002 | | 1.002 | | | |
| n_eff discrimination | 1091 | 2413 | | 2132 | | | |
| **Exp 2** | | | | | | | |
| Rhat identification | 1.003 | 1.000 | | 1.001 | | | |
| n_eff identification | 1031 | 2552 | | 2457 | | | |
| Rhat discrimination | 1.000 | 1.000 | | 1.000 | | | |
| n_eff discrimination | 1718 | 2735 | | 2572 | | | |
| **Exp 3** | | | | | | | |
| Rhat identification | 1.009 | 1.001 | | 1.002 | | | |
| n_eff identification | 725 | 2319 | | 1893 | | | |
| Rhat discrimination | 1.002 | 0.999 | | 1.000 | | | |
| n_eff discrimination | 1683 | 3562 | | 3431 | | | |
| **Exps 4-7 full model** | Target duration | congruency | Cue type | Dur x congr | Congr x cue | Dur x cue | triple |
| Rhat discrimination | 1.001 | 1.004 | 1.002 | 1.003 | 1.003 | 1.001 | 1.004 |
| n_eff discrimination | 1326 | 1832 | 1300 | 2073 | 1943 | 1208 | 2042 |

Indications of a good quality of fit are: Rhat values nearing 1.0 indicate that independent parameter estimates have converged to the same value. Rhat < 1.01 are usually considered sufficient for appropriate convergence[4]. Effective sample size (n_eff) above 1000 indicate stable estimates of parameter values[5].

#### SI References


1. Debrenne, M. Le dictionnaire des associations verbales du français et ses applications. *Variétés, variations and formes du français. Palaiseau: Éditions de l'Ecole polytechnique* 355–66 (2011).
2. Ferrand, L. & Alario, F.-X. Normes d'associations verbales pour 366 noms d'objets concrets. *L'Année psychologique* **98**, 659–709 (1998).
3. Ferrand, L. Normes d'associations verbales pour 260 mots «abstraits». *L'Année psychologique* **101**, 683–721 (2001).
4. Vehtari, A., Gelman, A., Simpson, D., Carpenter, B. & Bürkner, P.-C. Rank-Normalization, Folding, and Localization: An Improved Rˆ for Assessing Convergence of MCMC (with Discussion). *Bayesian Analysis* **16**, 667–718 (2021).
5. Bürkner, P.-C. brms: An R package for Bayesian multilevel models using Stan. *Journal of statistical software* **80**, 1–28 (2017).
